# A panel of KSHV mutants in the polycistronic kaposin locus for precise analysis of individual protein products

**DOI:** 10.1101/2021.06.11.448153

**Authors:** Mariel Kleer, Grant MacNeil, Eric S. Pringle, Jennifer A. Corcoran

## Abstract

Kaposi’s sarcoma-associated herpesvirus (KSHV) is the cause of several human cancers including the endothelial cell (EC) malignancy, Kaposi’s sarcoma. Unique KSHV genes absent from other human herpesvirus genomes, the “K-genes”, are important for KSHV replication and pathogenesis. Among these, the kaposin transcript is highly expressed in all phases of infection, but its complex polycistronic nature has hindered functional analysis to date. At least three proteins are produced from the kaposin transcript: Kaposin A (KapA), B (KapB), and C (KapC). To determine the relative contributions of kaposin proteins during KSHV infection, we created a collection of mutant viruses unable to produce kaposin proteins individually or in combination. Kaposin-deficient latent iSLK cell lines displayed reduced viral genome copy number and often exhibited small LANA nuclear bodies; despite this, all were capable of progeny virion production. Primary infection with ΔKapB virus revealed decreased LANA expression and viral genome copy number, yet providing KapB protein *in trans* failed to complement these defects, suggesting a requirement for the *kaposin* locus *in cis*. Our previous work showed that KapB was sufficient to recapitulate the elevated proinflammatory cytokine transcripts associated with KS via the disassembly of RNA granules called processing bodies (PBs). We now show that KapB is necessary for PB disassembly during latent KSHV infection. These findings demonstrate that our panel of kaposin-deficient viruses enables precise analysis of the respective contributions of individual kaposin proteins to KSHV replication. Moreover, our mutagenesis approach serves as a guide for the functional analysis of other complex multicistronic viral loci.

**Importance:** Kaposi’s sarcoma-associated herpesvirus (KSHV) expresses high levels of the kaposin transcript during both latent and lytic phases of replication. Due to its repetitive, GC-rich nature and polycistronic coding capacity, until now no reagents existed to permit a methodical analysis of the role of individual kaposin proteins in KSHV replication. We report the creation of a panel of recombinant viruses and matched producer cell lines that delete kaposin proteins individually or in combination. We demonstrate the utility of this panel by confirming the requirement of one kaposin translation product to a key KSHV latency phenotype. This study describes a new panel of molecular tools for the KSHV field to enable precise analysis of the roles of individual kaposin proteins during KSHV infection.

## Introduction

Kaposi’s sarcoma-associated herpesvirus (KSHV) is the infectious cause of the endothelial cell (EC) neoplasm, Kaposi’s sarcoma (KS) and two rare lymphoproliferative disorders: primary effusion lymphoma (PEL) and multicentric Castleman disease (MCD) (1–3). Like all herpesviruses, KSHV establishes persistent, life-long infection of its human host, and displays two modes of infection, latent and lytic replication. Latency is the default replication program upon *de novo* infection in most cell types (4, 5). Following a transient period of lytic gene expression which serves to amplify genome copy number and evade the intrinsic immune response (6, 7), the viral episome is circularized and tethered to the host chromosome by the viral latency-associated nuclear antigen (LANA) (8). This results in formation of microscopically visible LANA-nuclear bodies (NB), which correlate with intracellular viral genome copy number (9–13). In the latent state, the viral genome is passively replicated and unevenly partitioned to daughter cells by host cell machinery (10, 14, 15). The viral genome will remain in a highly chromatinized state until expression of the viral lytic switch protein, replication and transcription activator (RTA), which is both necessary and sufficient for lytic reactivation (16–18). Following reactivation, lytic gene expression follows a prescribed temporal cascade with genome replication marking the transition from early to late gene expression (19, 20). More recent analyses using single cell approaches show that lytic reactivation is quite heterogenous in terms of viral gene expression, host cell responses and outcomes (13, 21–23). The lytic replication phase produces progeny virions enabling transmission of the virus and culminates in cell death.

The KSHV-infected cells in KS lesions are predominantly proliferating endothelial cells (ECs) with an abnormal elongated or ‘spindled’ morphology. The majority of these tumor ECs exhibit latent KSHV infection, whereas lytic replication is limited (24–27). Lytic replication is hypothesized to play a role in KS, likely due to the ongoing production of progeny virions as well as the release of inflammatory and angiogenic factors (14, 28, 29). Consistent with this, limiting viral lytic replication caused KS regression, suggesting that spontaneous lytic reactivation is required for ongoing infection of naïve cells in the tumor environment and supports the cancer (30–32).

As latently infected cells comprise the bulk of the KS lesion, the contribution of KSHV latent gene expression to tumorigenesis has been explored extensively in both animal and cell culture models (29, 33–37)*. In vitro* infection of primary ECs with KSHV recapitulates many of the features of KS tumors, including efficient establishment of viral latency. During latency, gene expression is limited to six consensus protein products produced from an approximately 10kbp region of the viral genome termed the latency locus (LANA, viral cyclin [v-Cyclin], and viral fas-associated death domain (FADD)-like interleukin-1-β-converting enzyme (FLICE) inhibitory protein [v-FLIP], Kaposins [Kap] A, B, and C) and 12 pre-miRNAs that are processed into at least 25 mature miRNAs (38–41) Using ectopic expression models, several latent gene products have been shown to contribute to the establishment and maintenance of viral latency as well as phenotypes associated with KS tumors (reviewed in (42, 43), but these studies have predominantly focused on LANA, v-Cyclin and v-FLIP while the contributions of the polycistronic kaposin locus is less clear. The kaposin mRNA was first identified as a marker of KSHV latent infection in KS tumours in 1997 (27), and it remains the most abundant viral transcript in KS tumor isolates (44). The kaposin locus comprises a significant fraction of KSHV coding capacity during latency and kaposin transcription is also upregulated during lytic replication, suggesting that this region of the viral genome is important (45). Despite this, we still know very little about the role of the kaposin locus, or the proteins it encodes, during viral replication.

The kaposin transcript is polycistronic and can be translated into at least three polypeptides, KapA, KapB and KapC (Figure 1A) (45), though we and others have observed multiple banding patterns on immunoblots that suggest additional translation products may also be derived from this locus (45, 46). Translation of KapA is initiated at a canonical AUG start codon located distal to the 5’ end of the transcript and encodes a small membrane spanning protein (47–49). KapC is translated in the same reading frame as KapA, making its carboxy-terminal region identical to KapA. However, translation of KapC is initiated at one of two non-canonical CUG start codons proximal to the 5’ end of the transcript. Therefore, in addition to the KapA ORF, the KapC translation product contains two 23-amino acid direct repeat (DR) regions termed DR1 and DR2 (DR1: PSSPGTWCPPPREPGALLPGNLV and DR2: APQEPGAAHPRNPARRTPGTRRG) which are derived from three sets of GC-rich 23-nucleotide repeats (Figure 1A-B). Translation of KapB is initiated at one of two possible non-canonical CUG or GUG start codons that are most proximal to the 5’ end of the kaposin mRNA, but uses an alternative reading frame to that of KapA/C. The resulting polypeptide is comprised largely of the same two 23-amino acid DRs (DR1 and DR2) found in KapC (Figure 1B) but lacks the C-terminal membrane spanning domain found in both KapA and KapC. Moreover, different isolates of KSHV display expansion or contraction of both direct repeats, the functional significance of which is unknown, yet regardless of the diversity in size, the core DR1 and DR2 repeats remain constant (45).

**Figure 1.**
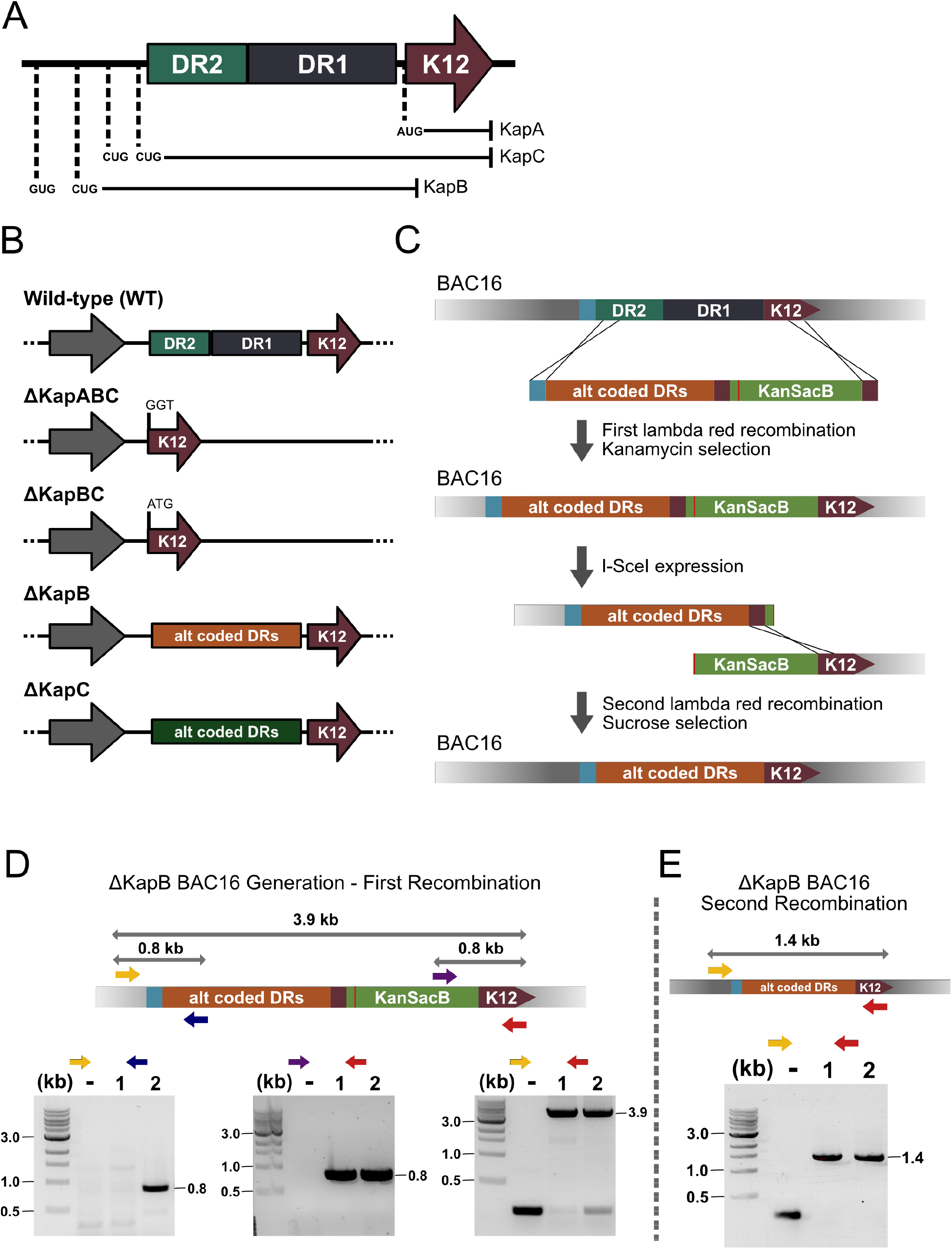
Lambda Red recombination facilitates the creation of five kaposin-deficient BAC16 DNA constructs. **(A)** A simplified version of the kaposin transcript is depicted in the 5’ to 3’ orientation. Open reading frames for kaposin A (KapA), kaposin B (KapB), and kaposin C (KapC) are indicated with corresponding AUG, GUG and CUG start codons for each. DR1 = direct repeat 1, DR2 = direct repeat 2, K12 = the KapA ORF. **(B)** An example of the re-coding strategy used to create the BAC16ΔKapB virus construct is shown. Wildtype nucleotide and corresponding amino acid repeats are show above with recoded nucleotides (red) and corresponding amino acid changes depicted below. **(C)** The Lambda Red recombination method used to create all kaposin-deficient BAC16 viruses is shown; the strategy for BAC16ΔKapB is depicted as an example. **(D)** Each kaposin-deficient BAC16 construct is shown relative to that of the wildtype *kaposin* locus in its genomic position (only reverse strand is shown for simplicity).

Our knowledge about individual kaposin protein function is derived largely from ectopic expression of individual kaposin proteins. KapA was reported to promote cell transformation *in vitro* and *in vivo* (47, 48); however, more recent work showed that this function stems from one of the viral miRNAs embedded within the KapA ORF (50). Previous work from our group has shown that KapB is sufficient to induce angiogenesis in an *in vitro* EC tubule formation assay and to promote actin stress fibers and elongation of ECs into a spindled morphology that recapitulates the morphology of latently infected KS tumor cells (51). KapB also is sufficient to induce the disassembly of RNA granules called processing bodies (PBs) (51, 52). PBs are important sites for the constitutive turnover or translational suppression of inflammatory cytokine mRNAs; PB loss correlates with enhanced levels of inflammatory cytokine transcripts that are normally targeted to PBs via an AU-rich destabilizing element (51, 53–57). However, our previous studies could not determine if KapB alone is necessary for these phenotypes during KSHV latent infection as we used RNA silencing to knockdown the kaposin transcript, a strategy that reduced expression of all kaposin proteins (51). No other studies have determined the role of individual kaposin translation products in KSHV latent or lytic replication cycles.

Here, we describe our use of the KSHV BAC16 bacterial artificial chromosome (BAC) and lambda Red recombination system (58) to construct a panel of viruses defective for specific *kaposin* locus products. We have constructed a triple deletion BAC16ΔKapABC, a double deletion BAC16ΔKapBC, and two single deletions BAC16ΔKapB and BAC16ΔKapC (Figure 1C). We confirmed that these viruses no longer express the deleted kaposin-derived proteins and show that all kaposin-deficient viruses are competent for genome replication and production of progeny virions. However, we show that ΔKapB is significantly impaired, with markedly reduced genome copy number, small LANA nuclear bodies, and lower LANA expression. Moreover, these defects could not be complemented by providing KapB *in trans*. Additionally, ΔKapB fails to cause PB disassembly following *de novo* infection of primary ECs, showing for the first time that KapB is necessary for PB disassembly during KSHV latency. With this panel of kaposin recombinant KSHV viruses, it is now feasible to interrogate, with unprecedented precision, the role of individual kaposin protein products in the context of KSHV latent or lytic infection.

## Materials and Methods

### Cell culture

All cells were grown at 37°C with 5% CO_2_ and atmospheric O_2._ HEK293T cells (ATCC) were cultured in DMEM (Thermo Fisher) supplemented with 100 U/mL penicillin, 100 µg/mL streptomycin, 2 mM L-glutamine (Thermo Fisher) and 10% fetal bovine serum (FBS; Thermo Fisher). iSLK.RTA cells (59) were cultured likewise with the addition of 1 µg/mL puromycin and 250 µg/mL geneticin (Thermo Fisher). BAC16-iSLK.RTA cells were cultured similar to iSLK.RTA cells, with the addition of 1200 µg/mL hygromycin B (Thermo Fisher). Human umbilical vein endothelial cells (HUVECs, Lonza) were cultured in endothelial cell growth medium (EGM-2) (Lonza). HUVECs were seeded onto gelatin (0.1% w/v in PBS)-coated tissue culture plates or glass coverslips.

### Lambda Red recombination for BAC16 mutagenesis

All mutagenesis was carried out in the BAC16 background (58) supplied in the GS1783 strain of *Escherichia coli (E.coli)* (A generous gift from Dr. Jae Jung). All restriction enzymes were purchased from New England Biolabs (NEB) and all primers were purchased from Invitrogen.

A kanamycin resistance and levansucrase enzyme cassette (KanSacB) was amplified using primers (Table 1) with 40bp overhanging sequences that correspond to the K12 sequence of the *kaposin* locus in the BAC16 genome. These primers also allowed for the introduction of BamHI and NotI restriction endonuclease sites. The resulting PCR product was then subcloned into pcDNA3.1+ (Addgene) using BamHI and NotI restriction endonucleases (pcDNA3.1+-KanSacB).

**Table 1:**
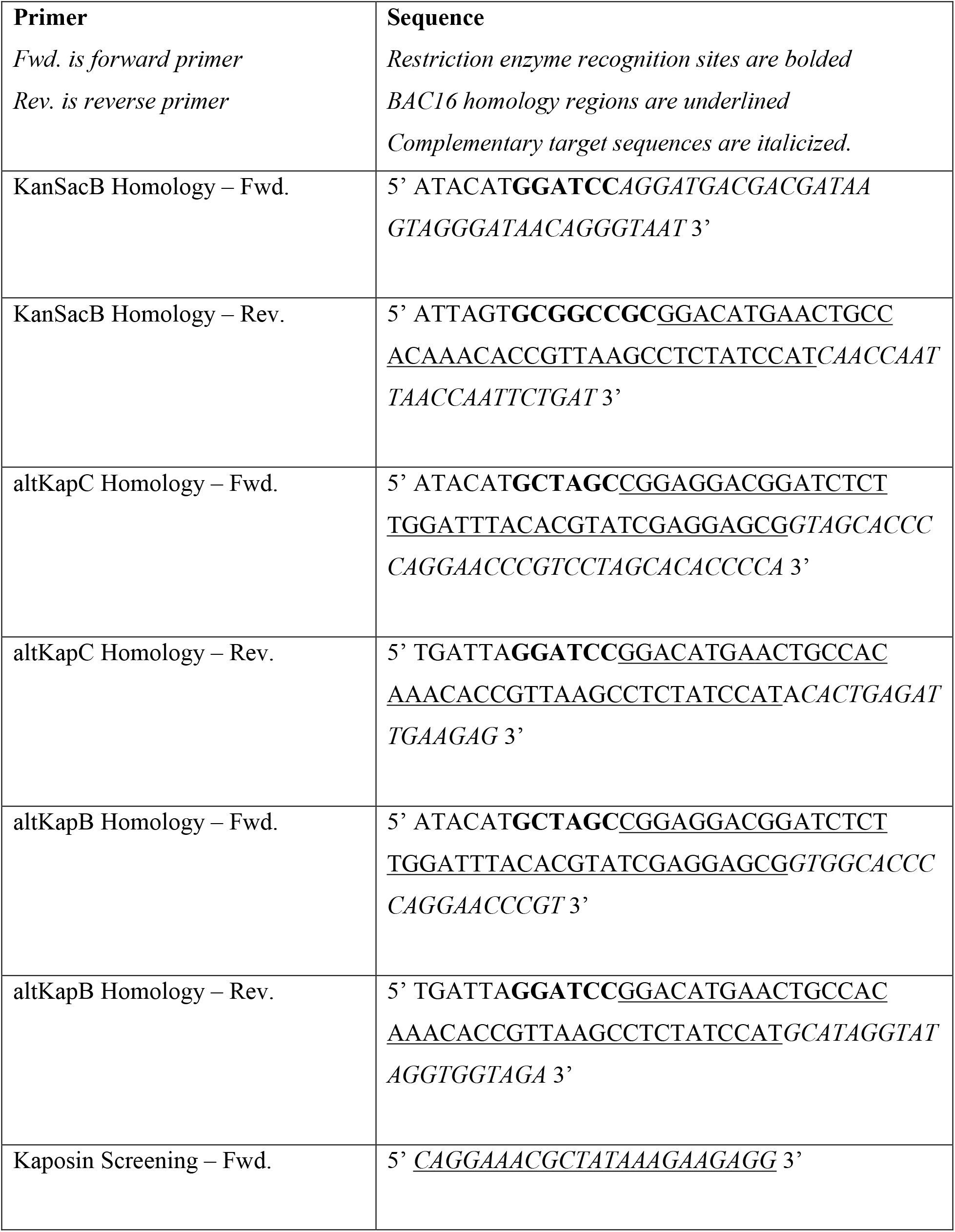
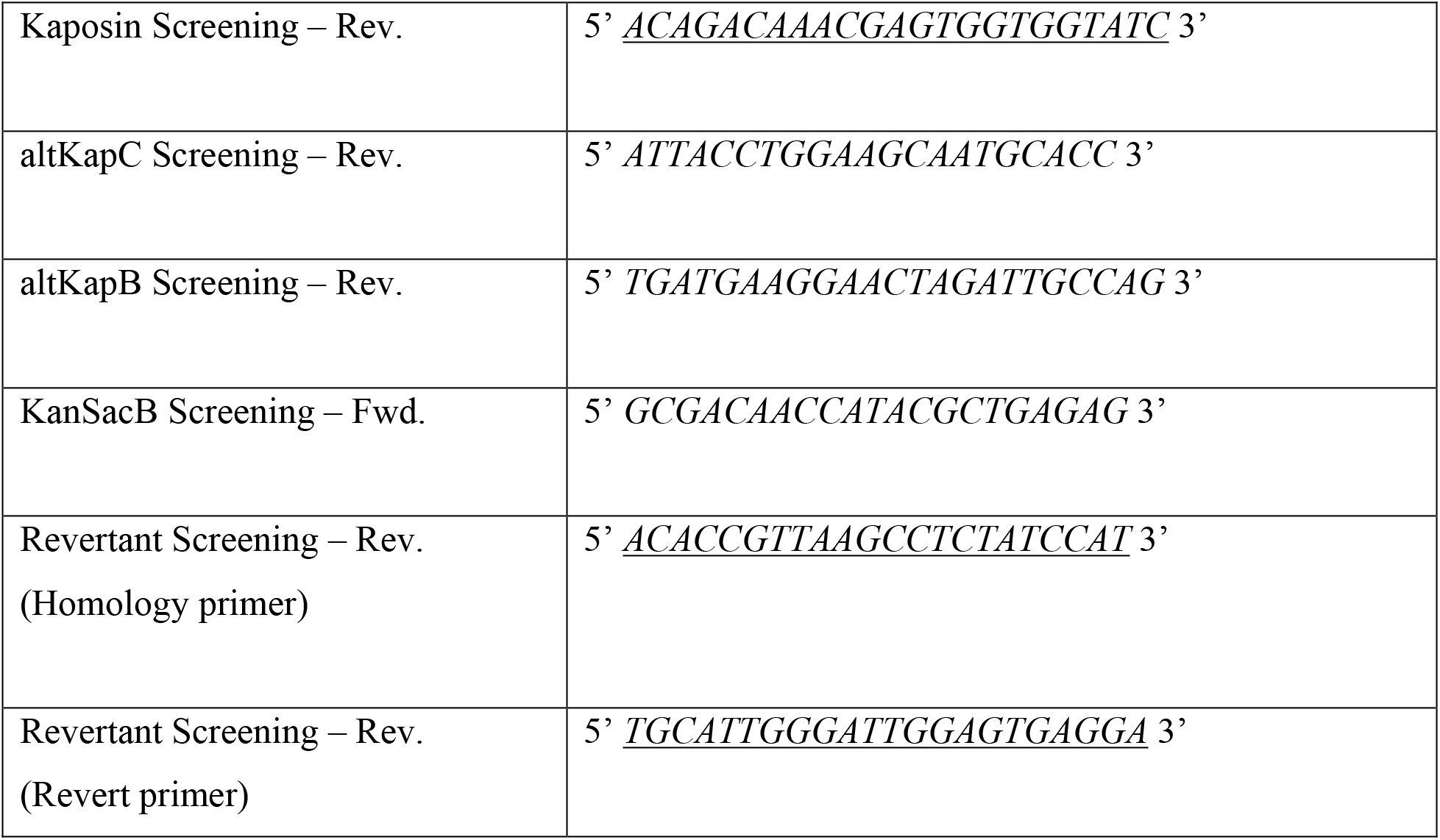
BAC16 Mutagenesis Primers.

To create single deletions of KapB and KapC, two direct repeat (DR) region gene blocks were synthesized (BioBasic) where the third nucleotide of each codon of DR1 and DR2 was altered to insert premature stop codons and nonsynonymous mutations into one frame, while creating synonymous mutations in the other. This allowed the primary amino acid sequence of either KapB or KapC to be maintained while eliminating expression from the other ORF and vice versa. Primers were used to amplify the gene block with 40bp overhanging sequences that: 1) flank the gene block, 2) are homologous to the upstream and downstream regions of the DRs in the BAC16 genome, and 3) introduce NheI and BamHI restriction sites. The resulting PCR product was then subcloned into pcDNA3.1+-KanSacB (described above) using NheI and BamHI restriction sites (pcDNA3.1+-geneblock-KanSacB).

Lambda Red recombination was conducted in the GS1783 *E.coli* strain, which expresses the I-Sce enzyme and, in a temperature (42°C) dependent manner, the lambda Red recombinase genes (beta, exo and gam). BAC16-GS1783 *E. coli* were cultured at 30°C in Lysogeny Broth (LB) containing 30 *μ*g/mL chloramphenicol (Sigma-Aldrich) until reaching an optical density (OD_600_) of 0.6-0.8. Cultures were incubated for 30 minutes at 42°C prior to pelleting at room temperature at 3000 x *g* for 5 minutes. Bacterial pellets were resuspended in ddH_2_0 and pelleted again at 3000 x *g* for 5 minutes. This step was repeated once more, following which bacterial pellets were resuspended in 50 *μ*L ddH_2_0 and placed on ice. Meanwhile, pcDNA3.1+-geneblock-KanSacB was digested with NheI and NotI restriction endonucleases to generate a linear DNA fragment for recombination. This was purified by 1% agarose gel electrophoresis and gel extracted according to QIAquick gel extraction kit (QIAGEN) standard protocol. The purified fragment was electroporated (2.5kV, 25*μ*F BioRad Gene Pulser®II) into 50 *μ*L of ddH_2_O-washed BAC16-GS1783 *E.coli*. Bacteria was recovered for 1 hour in 1 mL of LB media at 30°C, plated onto LB agar plates containing 50 µg/mL kanamycin, and incubated at 30°C overnight (O/N). Colonies were screened for successful recombination using colony PCR and two sets of screening primers designed to assess insert size and orientation (Table 1).

Colonies that produced bands of the expected size with both primer sets were inoculated into 1 ml of LB containing 1% arabinose (Sigma-Aldrich) for 1 hour at 30°C to induce I-Sce expression to linearize the inserted KanSacB cassette. Bacteria were then incubated at 42°C for 30 minutes to induce recombinase gene expression and recovered for 1 hour at 30°C prior to plating on LB agar plates containing 30 *μ*g/mL chloramphenicol and 5% sucrose, which selects for the loss of SacB that synthesizes a toxic compound in the presence of sucrose. Positive colonies were replica-plated on LB agar plates containing 50 µg/mL kanamycin. Those colonies that grew on both chloramphenicol and sucrose containing plates, and which were no longer kanamycin resistant, were screened by colony PCR for having undergone the second recombination to remove the KanSacB cassette and colony PCR products were verified by Sanger sequencing using primers as listed (Table 1).

### Isolation of BACmid DNA

BACmid DNA was isolated from GS1783 *E.coli* cultures of 5 mL of LB containing 30 *μ*g/mL chloramphenicol grown O/N at 30°C. Bacteria was pelleted at 3000 x *g* for 5 minutes, re-suspended in buffer P1 containing RNaseA, lysed using alkaline lysis in buffer P2 and neutralized with buffer N3 (All reagents from QIAprep kits, QIAGEN). Following addition of buffer N3, lysates were placed on ice for 10 minutes prior to centrifugation at 17 000 x *g* for 10 minutes to pellet precipitate. 750 µL of supernatant was then mixed with 750 µL of 100% isopropanol and placed on ice for 10 minutes prior to centrifugation at 17 000 x *g* for 10 minutes to pellet DNA. Supernatants were removed and the DNA pellet was left to air dry before being resuspended in 50 µL 10mM Tris-Cl, pH 8.5. DNA was used immediately for transfection.

### Whole genome sequencing of KSHV BACs

BACmid DNA was isolated from GS1783 *E.coli* cultures of 400 mL LB containing 30 *μ*g/mL chloramphenicol grown O/N at 30°C. Bacteria was pelleted at 4000 x *g* for 10 minutes at 4°C. DNA was then extracted using the Qiagen Large Construct Kit in accordance with manufacturer’s protocol. Critically, this isolation includes an exonuclease step which removes sheared *E.coli* DNA thereby allowing greater coverage of the BAC16 genome.

Whole genome sequencing was performed by the Centre for Health Genomics and Informatics (University of Calgary, Calgary, AB). Briefly, libraries were prepared using a NEB ultra II DNA Library Prep Kit for Illumina (NEB), quantified using a Kapa qPCR Library Kit (Roche), and sequenced using a MiSeq 500 v2 nano run (MiSeq Reagent Kit v2, Illumina) to generate 250+250bp paired end sequences. Raw reads were imported into Geneious Prime (60), paired and trimmed according to default settings. Reads were then aligned to the BAC16 reference genome (NCBI accession #GQ994935). Average coverage was between ∼670 and 1400x. Variations were identified based on a minimum coverage of 5 and minimum frequency in variation of 0.25.

### Restriction length fragment polymorphism

Isolated BACmid DNA was digested overnight using NheI restriction enzyme (New England Biolabs) at 37°C. Digested DNA fragments were then subjected to 1% agarose pulsed-field gel electrophoresis (PFGE) (Bio-Rad) using 0.5% Tris/Borate/EDTA (TBE) buffer and the following conditions: 6 V/cm, 120° field angle, and switch time linearly ramped from 1 s to 5 s over 14-15 h (CHEF DR III, Bio-Rad). DNA bands were visualized by staining the gel with a solution of 0.5 µg/mL ethidium bromide; excess stain was washed away with deionized water and the gel was imaged on a ChemiDoc Touch Imaging System (Bio-Rad). DNA molecular weight standard (BioRad, CHEF DNA size 8-48-kb) was used as reference.

### Generation of Stable BAC16-iSLK.RTA cell lines

6-well plates of 70% confluent 293T cells in antibiotic-free 10% FBS DMEM were washed once with PBS, and media was replaced with 1 mL serum-free, antibiotic-free DMEM per well prior to transfection. 5 uL of re-suspended BAC16 DNA was incubated with 100 uL Opti-MEM reduced serum media (Thermo Fisher) for 5 minutes. Separately, 18 uL of polyethylenimine (PEI, Polysciences) was incubated with 100 uL Opti-MEM, also for 5 minutes. These two mixtures were combined, incubated for 15 minutes, and added dropwise to each well. Cells were incubated at 37°C for 4 hours at which time the media was replaced with antibiotic-free 10% FBS DMEM before further incubation at 37C. This transfection procedure was used to create latent 293T stable cell lines shown in Figure 3.

At 48 hours post transfection, cells were expanded into 10-cm dishes. One day later, media was removed and replaced with fresh antibiotic-free 10% FBS DMEM containing 20 ug/mL hygromycin (Thermo Fisher Scientific). The concentration of hygromycin was steadily increased until a final concentration of 100 *μ*g/mL was reached. When a confluent monolayer of hygromycin-resistant BAC16 293T cells was achieved, cells were split once prior to liquid N2 storage in 90% FBS and 10% dimethyl sulfoxide (DMSO, Sigma-Aldrich).

To generate latent iSLK cells, each population of BAC16-transfected 293T cells was individually co-cultured with puromycin-resistant iSLK.RTA cells in a 6-well dish at a 2:1 ratio. Once cells reached 80% confluency, media was replaced with antibiotic-free 10% FBS DMEM containing 1 mM sodium butyrate (NaB, Sigma-Aldrich) and 20 ng/mL 12-O-tetradecanoylphorbol-13-acetate (TPA, Sigma-Aldrich) to induce latent BAC16-293T cells to produce progeny virions. At 96 hours post reactivation, cells were expanded into 10 cm dishes. One day following expansion, media was replaced with antibiotic-free 10% FBS DMEM containing 10 ug/mL puromycin and 100 *μ*g/mL hygromycin. Hygromycin concentration was steadily increased in approximately 50 *μ*g/mL increments until a final concentration of 240 *μ*g/mL was reached. Once a near-confluent monolayer of BAC16-iSLK.RTA cells was achieved, cells were split once prior to liquid N_2_ storage as above. This procedure was used to create the latent iSLK cell lines shown in Figure 3. A single latent iSLK cell line was created for ΔKapABC, ΔKapBC and ΔKapC, while we derived four latent iSLK cell lines for ΔKapB independently, as shown in Figure 6.

### Production of KSHV BAC16 Wild-type and Recombinant Virus

Each population of BAC16-iSLK cells were reactivated using 1 ug/mL Doxycycline (Dox, Sigma-Aldrich) and 1 mM NaB in antibiotic-free 10% FBS DMEM. At 72 or 96 hours post infection, supernatants were collected and clarified at 4000 x *g* to remove cellular debris and virus-containing supernatant stored at −80°C prior to use.

### *De novo* infection of Naïve HEK293T of HUVEC cells

Sub-confluent (50-70%) 12-well plates of 293T or HUVEC cells were incubated with viral inoculum diluted in antibiotic and serum free DMEM containing 8 *μ*g/mL hexadimethrine bromide (polybrene; Sigma-Aldrich). Plates were centrifuged at 800 x *g* for 2 hours at room temperature (61), the inoculum was removed and replaced with either antibiotic-free 10% FBS DMEM (293T) or full EGM-2 (HUVECs). Cells were then returned to the 37°C incubator.

### Immunofluorescence

Cells were seeded onto 18mm round (#1.5, Electron Microscopy Sciences) coverslips, infected or treated as noted for each experiment and fixed for 10 minutes at room temperature in 4% (v/v) paraformaldehyde (Electron Microscopy Sciences) in PBS. Samples were permeabilized with 0.1% (v/v) Triton X-100 (Sigma-Aldrich) in PBS for 10 min at room temperature and blocked in 1% human AB serum (Sigma-Aldrich) for 1 hour at room temperature. Primary antibody was diluted in 1% human AB serum (Sigma-Aldrich) in PBS at the concentrations in Table 2 and incubated overnight. The following day secondary antibody was likewise diluted in 1% human AB serum, alongside Hoechst (ThermoFisher Scientific) for nuclear staining, at the concentrations in Table 2 and incubated for one hour. Samples were mounted with Prolong Gold AntiFade mounting media (ThermoFisher Scientific). Images were captured using a Zeiss AxioObserver Z1 microscope with the 40X objective unless otherwise stated.

**Table 2:**
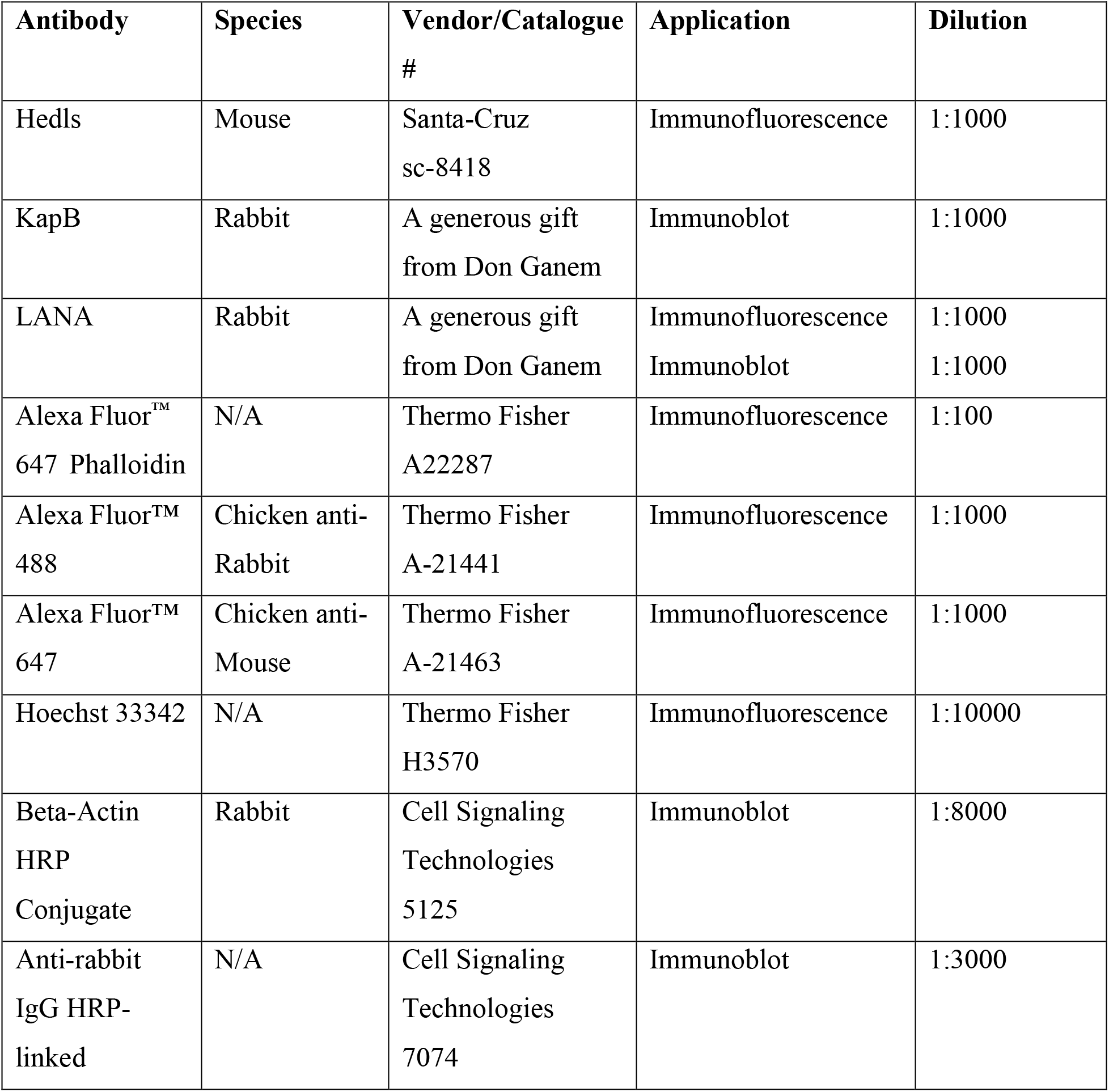
Antibodies.

### Image analysis

Image analysis was performed using CellProfiler (cellprofiler.org), an open source software for image analysis (62). Quantification of hedls/EDC4 and LANA-nuclear body (NB) puncta was performed as previously described (63) with the following modifications. Rather than staining cells with wheat germ agglutinin to identify cell periphery, individual cell cytoplasms were defined based on a 150-pixel propagation away from the Hoescht-stained nucleus. The number of Hedls/EDC4 puncta within a defined cell cytoplasm was reported. To quantify the number of puncta in KSHV-infected cells, a gating strategy was applied to include only cells with LANA-positive nuclei and defined cell cytoplasms for Hedls/EDC4 puncta quantification. LANA-NBs were defined by thresholding and size was measured using the “MeasureObjectSizeShape” function, respectively. Object area refers to the number of pixels within the identified region. At least 50 cells were quantified for each condition.

### Immunoblotting

Cells were lysed in 1x Laemmli buffer (4% SDS, 20% glycerol, 120 mM Tris-Cl (pH 6.8) and ddH_2_0) and stored at −20°C until use. The DC Protein Assay (Bio-Rad) was used to quantify total protein concentration as per the manufacturer’s instructions. 10-15 µg of protein lysate was resolved by SDS-PAGE on TGX Stain-Free acrylamide gels (BioRad). Membranes were blocked in 5% BSA or 5% skim milk in Tris buffered saline-Tween20 (TBS-T). Primary and secondary antibodies were diluted in 2.5% BSA or 2.5% skim milk, dilutions can be found in Table 1. Membranes were visualized using ProtoGlow ECL (National Diagnostics) and the ChemiDoc Touch Imaging system (BioRad).

### Intracellular viral genome qPCR

Confluent 6-well plates of BAC16-iSLK.RTA cells were lysed using a DNeasy Blood and Tissue Kit (Qiagen) according to manufacturer’s protocol. Isolated DNA was stored at −20°C until further use. DNA was diluted 1:30 prior to qPCR amplification using SsoFast EvaGreen Supermix (Bio-Rad) and viral ORF26 or cellular β-actin (control) primers as listed in Table 3. Fold change in viral genome copy number was determined using the ΔΔ-Quantitation cycle (Cq) method.

**Table 3:**
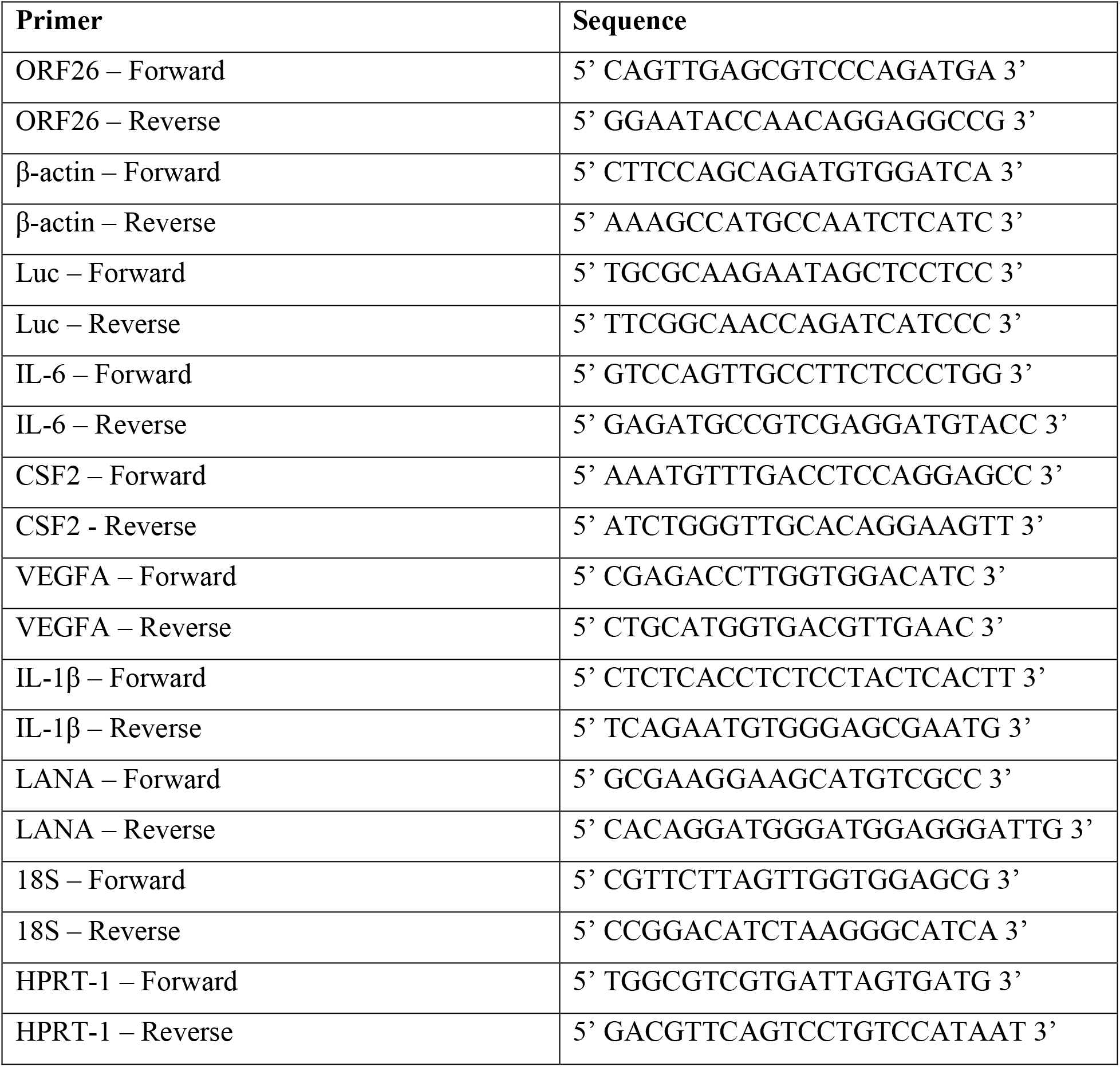
qPCR primers.

### DNase-protected viral genome qPCR

Supernatants from reactivated iSLK cells were harvested and clarified at 4000 x *g* for 5 minutes. Aliquots of 180 µL were stored separately at −80°C prior to use. Supernatants were thawed at 37°C and mixed with 20 µL of a 500U DNaseI (NEB), 10X buffer and nuclease free water mixture and incubated at 37°C for 30 minutes. DNA was then extracted using a DNeasy Blood and Tissue Kit with the following adjustments: per sample, 500 µg of salmon sperm DNA (Invitrogen) and 1 ng of a luciferase (*luc2*) containing plasmid (pGL4.26, Clontech) was added to buffer AL prior to lysis. DNA was not diluted prior to qPCR amplification using SsoFast EvaGreen Supermix (Bio-Rad) and viral ORF26 and *luc2* (control) primers as listed in Table 3. Since every viral particle contains one genome, and one genome contains a single copy of the ORF26 gene, the number of viral DNA molecules could be calculated. Fold change in viral DNase-protected genomes was determined using the ΔΔ-Quantitation cycle (Cq) method.

### Lentivirus Production and Transduction

All lentiviruses were generated using a second-generation system. Briefly, HEK293T cells were transfected with pSPAX2, MD2G, and the plasmid containing a gene of interest using polyethylenimine (PEI, Polysciences). psPAX2 was a gift from Didier Trono (Addgene plasmid #12260) and pMD2.G was a gift from Didier Trono (Addgene plasmid #12259). Viral supernatants were harvested 48 h post-transfection, clarified using a 0.45 µm polyethersulfone filter (VWR), and frozen at −80°C until use. For transduction, lentiviruses were thawed at 37°C and added to target cells in complete media containing 5 µg/mL polybrene (Sigma) for 24 h. The media was changed to selection media containing 5 µg/mL blasticidin (Thermo) and cells were selected for at least 48 h before proceeding with experiments.

### Transfection

12-well plates of 70% confluent 293T cells in antibiotic-free 10% FBS DMEM were washed once with PBS, and media was replaced with 1 mL serum-free, antibiotic-free DMEM per well prior to transfection. Cells transfected with 1 µg of plasmids containing a gene of interest (Table 4) using polyethylenimine (PEI, Polysciences). 4 hours post transfection, media was replaced with antibiotic-free 10% FBS DMEM.

**Table 4:** Plasmids.

| Plasmid | Use | Source | Mammalian Selection |
| --- | --- | --- | --- |
| pcDNA3.1 (+) | Empty Vector Control | Invitrogen V79020 | N/A |
| pcDNA3.1 KapB KS lung | Overexpression | C. McCormick (Dalhousie University) | N/A |
| pcDNA3.1 KapB BAC16 | Overexpression | Cloned from pUC57 KapB BAC16 (Bio Basic) | N/A |
| pLJM1 | Empty Vector Control | (105) | Blasticidin |
| pLJM1 KapB (pulmonary KS) | Overexpression | Cloned from pBMNIP-KapB (51) into pLJM1-BSD | Blasticidin |
| pMD2.G | Lentivirus generation | Addgene #12259 | N/A |
| psPAX2 | Lentivirus generation | Addgene #12260 | N/A |

### RT-qPCR

RNA was collected using a RNeasy Plus Mini Kit (Qiagen) according to the manufacturer’s instructions and stored at −80°C until further use. RNA concentration was determined using a NanoDrop One^C^ (Thermo) and 500 ng total was reverse transcribed using qScript XLT cDNA SuperMix (QuantaBio) using a combination of random and oligo (dT) primers according to the manufacturer’s instructions. cDNA was diluted 1:10 for all RT-qPCR experiments and SsoFast EvaGreen Mastermix (Biorad) was used to amplify cDNA. The ΔΔquantitation cycle (Cq) method was used to determine the fold change in expression of target transcripts. qPCR primer sequences can be found in Table 3.

### Statistics

Data shown are the mean ± standard deviation (SD). Statistical significance was determined using a paired Student’s *t*-test. *P* values are indicated in figures, (*, P < 0.05; **, P < 0.01; ***, P < 0.001; ****, P < 0.0001; ns, nonsignificant). All statistics were performed using GraphPad Prism version 9.0.1.

### Data Availability

All BAC16 virus sequences were deposited in the NCBI Sequence Read Archive (SRA) and are accessible through the following SRA accession numbers: SAMN19671530 (WT KSHV BAC16), SAMN19671876 (KSHV BAC16 ΔKapABC), SAMN19671877 (KSHV BAC16 ΔKapBC), SAMN19671902 (KSHV BAC16 ΔKapB), and SAMN19672118 (KSHV BAC16 ΔKapC).

## Results

### Construction of a panel of kaposin recombinant BAC16 KSHV viruses

Work on the *kaposin* locus has been limited by the lack of recombinant viruses that target this region. To address this limitation, we created a panel of recombinant viruses that delete KapA, KapB and KapC individually or in combination. These recombinant viruses were created using the KSHV bacterial artificial chromosome clone BAC16, which contains the pBelo45 plasmid inserted between the vIRF-1 and ORF57 genes of the JSC-1 KSHV isolate (58, 63). This insertion encodes a red fluorescent protein (RFP) and a hygromycin resistance gene under the control of a constitutive cellular promoter, to enable visual and chemical selection of BAC16-containing cells, respectively. The wildtype (WT) kaposin mRNA encodes three open reading frames (ORFs), KapA, KapB and KapC (Figure 1A). As a first step to create viruses that no longer made one or more of these protein products, we removed the DR regions that encompass the entire KapB ORF and most of the KapC ORF; in so doing, we created BAC16ΔKapBC. We then mutated the KapA start codon in the context of BAC16ΔKapBC to create BAC16ΔKapABC. However, creating single deletions of KapB and KapC by simple start codon mutagenesis proved to be extremely difficult due to the GC-rich and repetitive nature of this region. We decided to use an amino acid recoding strategy to construct BAC16 genomes that failed to code for either KapB or KapC, but not both. While both ORFs are comprised largely of 23-nucleotide DRs that translate into 23-amino acid DRs, the ORFs are in different reading frames. Using the BAC16 genome (accession #GQ994935), we designed two different gene blocks with the nucleotide sequence of the DRs recoded. The first DNA construct was composed of alternative codons that disrupted the protein coding capacity of the KapB ORF, due to several nonsynonymous or nonsense mutations in the DRs, but contained only synonymous nucleotide changes in the KapC ORF that did not disrupt its amino acid sequence (ΔKapB; Figure 1B). The second gene block coded for the inverse mutant; DRs were re-coded such that the KapC ORF contained nonsynonymous or nonsense mutations, while the KapB ORF contained synonymous mutations that did not disrupt its amino acid sequence (ΔKapC; Figure 1B). This strategy also drastically decreased the GC-content of DR region from 78% to 59.6% or 63.3% for ΔKapB and ΔKapC, respectively, facilitating the Lambda Red-based recombination events that followed and enabling us to use colony PCR screening during mutagenesis.

A two-step lambda Red-based recombination method in *E.coli* was used to facilitate BAC16 mutagenesis (Fig 1C) as in (64). We isolated each recombinant BACmid from *E.coli* and performed whole genome sequencing (WGS). Mapping of reads to the BAC16 reference genome (NCBI accession #GQ994935) using Geneious Prime (60) revealed an average coverage per nucleotide of between 670 and 1430x. Subsequent analysis of variants revealed that no off-target mutations had occurred during recombination (Figure 2A). Few variants were observed and these occurred within repetitive regions of the KSHV genome that also displayed low coverage, as previously shown (65–67). Outside of these, the only other variants detected were located in the region between the genes for vIRF-1 and ORF57 (Figure 2A). This is to be expected, as this is where the pBelo45 plasmid was inserted to facilitate the creation of BAC16, thereby altering the wild-type nucleotide sequence in this genomic location (58). We focused our attention to the kaposin DRs in WT BAC16. The repetitive nature of this sequence makes it difficult to resolve by WGS (68) and we observed low coverage. There was no coverage in the DR region for both ΔKapABC and ΔKapBC constructs, consistent with the complete removal of the DRs (Figure 2A). In contrast, we observed high read coverage of the recoded kaposin regions in ΔKapB and ΔKapC viruses as recoding decreased the repetitive and GC-rich nature of the DRs (Figure 2B, only WT and ΔKapB shown for simplicity). The consensus sequences produced for these regions matched our recoded gene blocks, reaffirming successful recombination. Given the difficulty in resolving DR regions through Illumina sequencing (68), we also checked the BAC16 genomes using restriction fragment length polymorphism. BAC16 DNA isolated from *E.coli* was digested using the restriction enzyme NheI, followed by pulse field gel electrophoresis (PFGE) to resolve fragments. We observed fragment sizes consistent with predictions from *in silico* analysis using NCBI accession #GQ994935 with the exception of the fragment comprised largely of terminal repeats (TRs). We predicted that the TR-containing fragment should migrate at 33 kbp; however, we did not observe a fragment of this size and instead observed an unexpected fragment migrating at approximately 18 kbp. This suggested a deletion of approximately 15 kbp, or 19 copies of the 801bp sequence, within this region. This deletion has previously been observed in BAC16 (69) because the highly repetitive nature of the TRs, as well as their high GC-content, renders them sensitive to recombination events (70, 71). The smaller TR fragment was found in WT BAC16 and in all five recombinant viruses, suggesting the event occurred prior to our mutagenesis, and that analysis of our panel kaposin-deficient BAC16 viruses relative to WT would be unaffected. PFGE analysis also revealed a second size discrepancy within the predicted kaposin-containing fragment which WGS was unable to resolve due to the repetitive nature of the region. Based on *in silico* analysis, we expected a 1200 bp fragment, yet the digestion of our WT BAC16 genome produced a 1000 bp fragment. This suggests that over time the WT BAC16 construct lost 200bp, or approximately three 23-amino acid repeats, within the DR region, making our recoded kaposin locus in ΔKapB and ΔKapC viruses longer than the kaposin locus in WT BAC16. Because different KSHV isolates possess variable numbers of repeats (45), it is not surprising to observe that this repetitive genomic area experienced DR contraction during BAC16 DNA copying, like has been observed for the *kaposin* locus in KSHV. In future studies, we hope to decipher whether there is any functional significance to the number of each kaposin DR1 and DR2 for KSHV replication.

**Figure 2.**
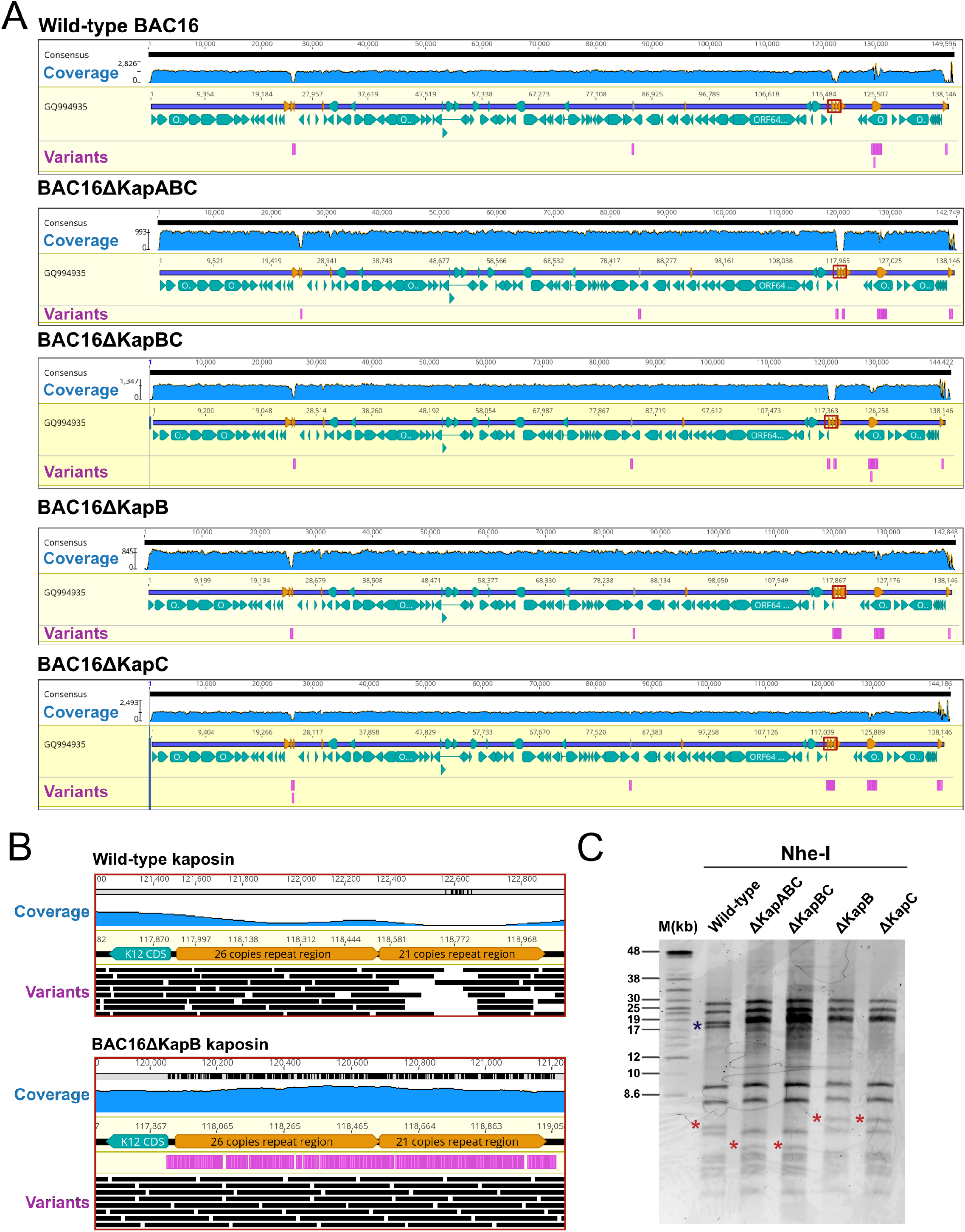
Confirmation of the panel BAC16 kaposin-deficient viruses. **(A)** BAC16 DNA was isolated from GS1783 *E.coli* harbouring recombinant or WT BAC16 DNA and then subjected to whole genome sequencing (WGS). Raw reads were paired, trimmed, and mapped to the BAC16 genome (NCBI accession #GQ994935) in Geneious Prime. Coverage for each recombinant is indicated in blue with an average coverage per nucleotide of between 670-1430x. BAC16 ORFs are depicted in turquoise, repeat regions in orange and the *kaposin* locus is outlined in red. Variants were detected based on a minimum coverage of 5 and minimum frequency variation of 0.25 and are depicted in magenta. **(B)** Representative zoom panels of read alignment to the *kaposin* locus of WT and ΔKapB are shown. **(C)** BAC16 DNA was isolated from GS1783 *E.coli* and digested using NheI and fragments were subjected to pulsed-field gel electrophoresis (PFGE). Based on *in silco* predictions, *kaposin* locus containing fragments are indicated with a red asterisk.

### Generation of stable BAC16-iSLK.RTA cell lines

After sequencing, the BAC16 mutant genomes were used to generate stable iSLK cell lines. A workhorse cell line for KSHV research, iSLK cells have been engineered to express the viral lytic switch protein, RTA, in a Doxycycline (Dox)-inducible manner (59). These cells are especially useful for KSHV research as they maintain tight control of latency and produce large quantities of virus following the addition of Dox (59). However, iSLK cells were refractory to transfection in our hands; thus, we first transfected BAC16 DNA into 293T cells to create stable BAC16-293Ts for each BACmid (Figure 3A), as in (69). The efficiency of 293T transfection was between 20-30%, as estimated by RFP expression. We gradually selected the transfected cells with hygromycin until a confluent monolayer of RFP+ 293T cells was achieved (Figure 3B). Via quantitative PCR (qPCR) for viral DNA (ORF 26), we analyzed intracellular genome copy number for each kaposin-deficient transfected 293T latent cell line and found that intracellular genome copy number was similar to WT in each case (Figure 3C).

**Figure 3.**
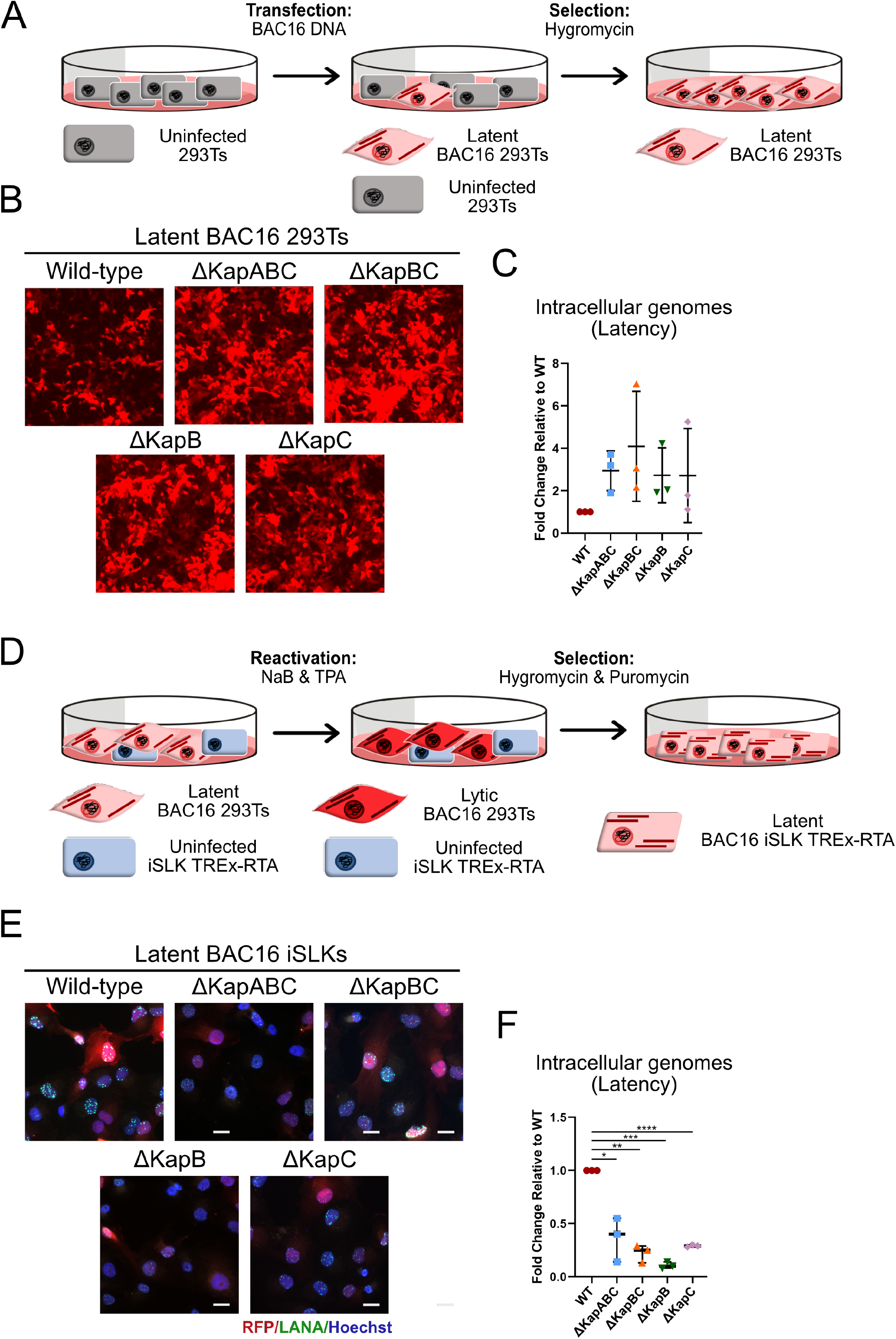
Generation of stable latent kaposin-deficient BAC16 iSLK cell lines by co-culture method. **(A-D)** 293T cells were transfected with recombinant or WT BAC16 DNA. 48 hours post transfection, cells were selected with hygromycin (increasing concentration from 50 μg/mL to 100 μg/mL) until a confluent layer of BAC16-containing 293T cells was achieved. **(B)** Images of BAC16-293Ts were captured using an EVOS-FL microscope with the 10X objective. **(C)** Latent BAC16-293T cells were trypsinzed, pelleted and lysed to extract DNA. qPCR was performed using ORF26 and β-actin specific primers. Values are represented as fold change relative WT. *n*=3; mean ± SD. **(D-F)** BAC16-293Ts were seeded in a 2:1 ratio with naïve iSLK cells. Cells were treated with 1 mM NaB and 20 ng/mL TPA for 96 hours, virus-containing supernatant was removed and replaced with media containing puromycin (10 μg/mL) and hygromycin (increasing concentration from 200 μg/mL to 1200 μg/mL) until a confluent layer of BAC16-containing iSLK cells was achieved **(E).** Latent iSLKs were fixed, permeabilized and immunostained using antisera for viral LANA (green) to visualize infected nuclei and Hoescht to visualize total cell nuclei (blue). Images were captured using Zeiss AxioObserver Z1 microscope with the 40X objective. Scale bar = 20 μm. **(F)** BAC16-containing iSLKs were trypsinized, pelleted and lysed to extract DNA. qPCR was performed using ORF26 and β-actin specific primers. Values are represented as fold change relative to WT. *n*=3; mean ± SD, (*, P < 0.05; **, P < 0.01; ***, P < 0.001; ****, P < 0.0001; ns, nonsignificant).

BAC16-containing 293T cells were then co-cultured with naïve iSLKs and reactivated using a histone deacetylase inhibitor (NaB) and a protein kinase C agonist (TPA) to produce progeny virions to enable primary infection of iSLKs (72–75). All kaposin-deficient viruses were able to infect naïve iSLKs judging by RFP expression. The population of kaposin-deficient latent iSLK cells for each recombinant was then expanded via gradual selection in the presence of hygromycin to eliminate uninfected cells and puromycin to eliminate BAC16-293T cells, until a confluent layer of RFP-positive cells was achieved (Figure 3D). Immediately after selection, we analyzed the features of our iSLK latent cell lines. RFP-positive iSLKs cells were then stained by immunofluorescence to visualize the viral latent protein LANA, which forms nuclear puncta called LANA nuclear bodies (NBs) during KSHV infection (9–12) that were visible in all our kaposin-deficient latent iSLK cell lines (Figure 3E), suggesting latency was successfully established. We observed that not all LANA-positive cells were RFP-positive, a discrepancy that has been reported by others (76). We also observed variable RFP intensity in our latent cell populations and an overall decreased intensity of RFP signal in kaposin-deficient cell lines relative to WT (Figure 3E). Using qPCR to quantify intracellular genome copy number of our panel of latent iSLKs, we revealed that kaposin-deficient iSLKs contained 10-40% of the genome copies observed in latent WT iSLKs (Figure 3F). These observations suggest the possibility that KSHV recombinant viruses that lack the *kaposin* locus display defects in their ability to establish latency and/or undergo genome amplification immediately following *de novo* infection, a step that serves to increase latent episome copy number (6, 7).

Considering the possibility that *kaposin*-deficient KSHV recombinant viruses could be less efficient at maintaining KSHV latency once already established, we decided to test how well each latent cell population would maintain the viral genome compared to WT BAC16 latent iSLKs in the absence of selective pressure. We passaged each latent cell line fifteen times in the absence of hygromycin and determined intracellular genome copy number at each passage. While WT BAC16 iSLKs displayed minimal episome loss over 15 passages (14, 77), we observed that two of our kaposin-deficient recombinant viruses displayed accelerated genome loss: ΔKapABC and ΔKapBC, whereas ΔKapB and ΔKapC genome copy number remained steady over time (Figure 4A-B). Taken together with Figure 3, these data suggest that the *kaposin* locus is important for the establishment of KSHV latency and may also contribute to its maintenance.

**Figure 4.**
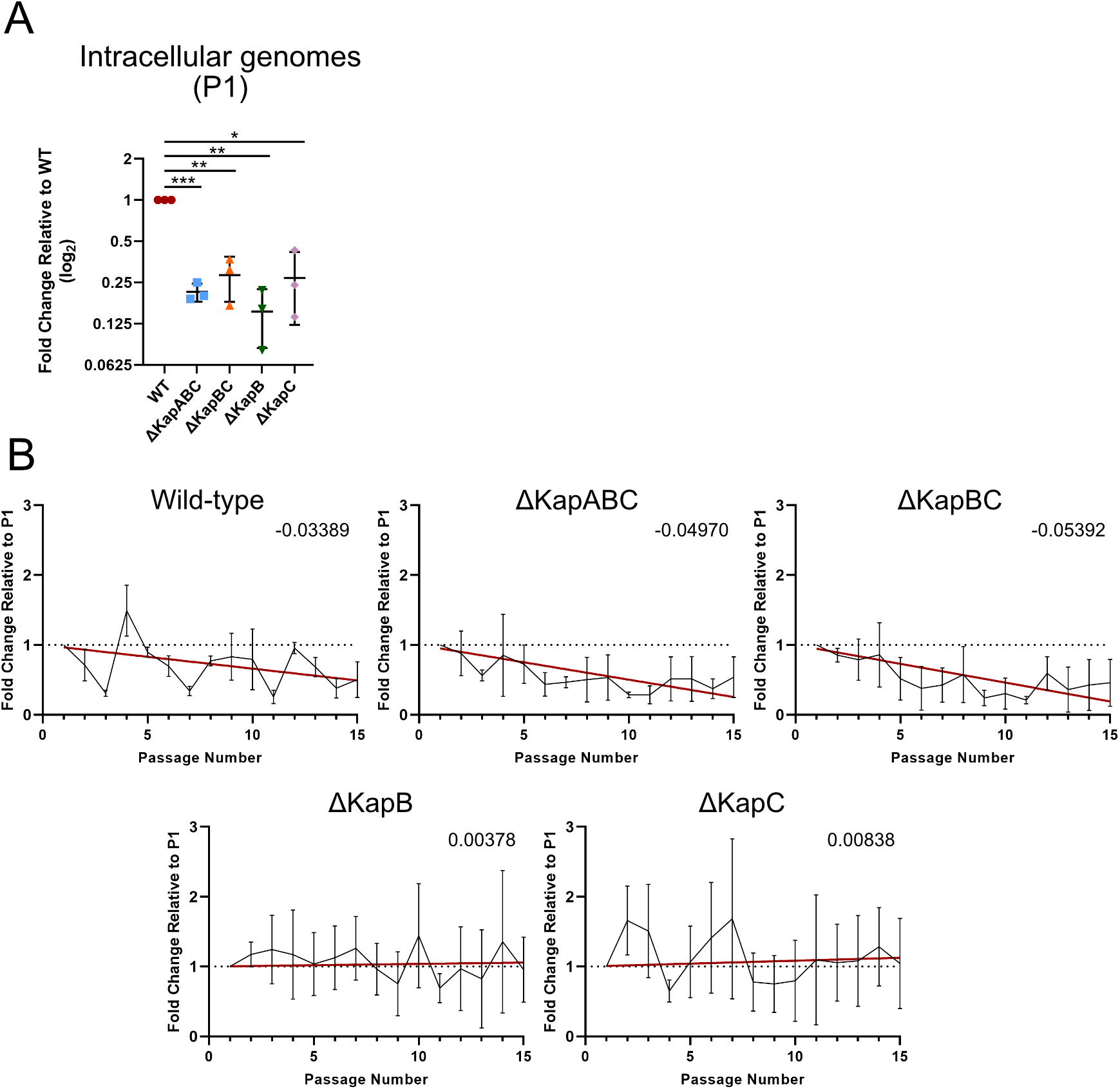
Removal of kaposin direct repeats accelerates KSHV episome loss from latent iSLK cells. Latent iSLK cells were serially passaged in the absence of hygromycin for a total of 14 passages. At each passage, a sample of cells was taken, pelleted and used for DNA extraction. To quantify levels of viral DNA, qPCR was performed using viral ORF26 and host β-actin specific primers. **(A)** Values are represented as fold change relative to genome copy number in wild-type (WT) BAC16 iSLK cells at passages 1 (P1) *n* = 3; means ± SD, (*, P < 0.05; **, P < 0.01; ***, P < 0.001; ****, P < 0.0001; ns, nonsignificant). **(B)** Values are represented as fold change relative to the corresponding starting passage (P1), for each recombinant virus over the course of the experiment. *n* = 3; mean ± SD. The red line was obtained using linear regression analysis and the slope of the line is depicted in the top right corner for each graph.

### Cells latently infected with kaposin-deficient viruses produce progeny virions when reactivated

Having created latently infected iSLK cell lines for each of our recombinant viruses, we first confirmed the absence of KapB and/or KapC protein products after 24 hours of reactivation of our ΔKapABC, ΔKapBC, and ΔKapB/ΔKapC latent iSLK cell lines by immunoblotting with a DR1-specific antibody (45) (Figure 5A). WT-infected cells exhibited a range of DR-specific banding as has been previously observed, suggesting additional isoforms of KapB or KapC can be translated (45, 46). Predominant bands at approximately 41kDa and 35kDa corresponded to the predicted molecular weights of full-length KapC and KapB, respectively. As expected, due to the absence of DR1 epitopes in these mutants, no KapB or KapC protein was detected in lysate from reactivated ΔKapABC or ΔKapBC iSLKs. We observed a faint band was detected at ∼46kDa in lysate derived from ΔKapB-infected iSLKs which corresponds to the predicted molecular weight of re-coded KapC (Figure 4A). This version of KapC is larger than that encoded by our WT BAC16 because of the loss of DRs in our version of WT BAC16 (Figure 2). A prominent band was consistently observed at ∼42kDa in lysate from ΔKapC-infected iSLKs which corresponds to the predicted molecular weight of re-coded KapB (Figure 5A), also larger than WT-derived KapB for the same reason (Figure 2). We attempted to produce an anti-KapA chicken yolk IgY antibody (78) using a customized antibody production service and an epitope predicted to be immunodominant; however, the antibody failed to detect KapA in WT infection or after ectopic expression and we did not use it further in these studies.

**Figure 5.**
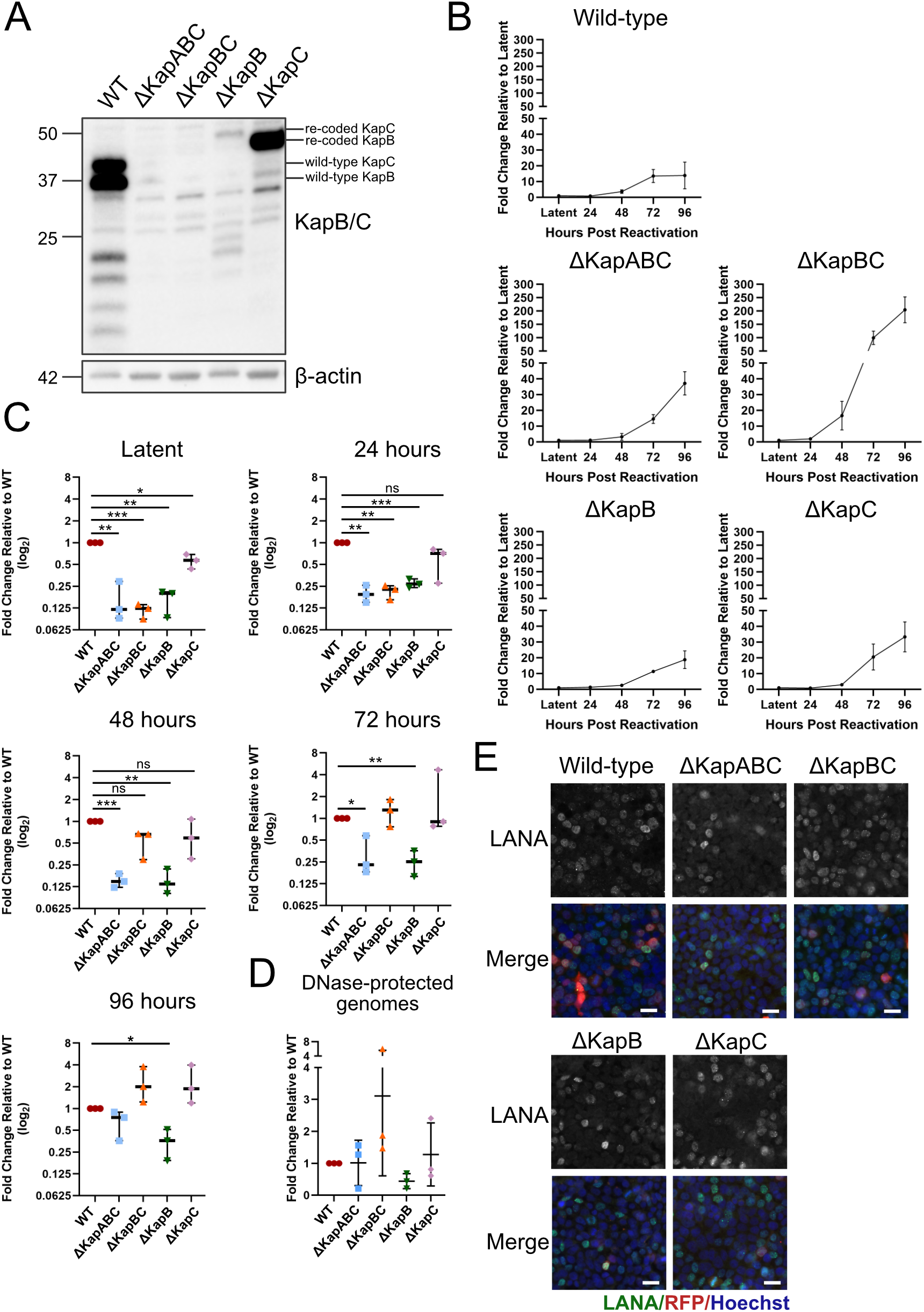
Kaposin-deficient BAC16 iSLK cells produce progeny virions when reactivated. **(A)** BAC16-iSLK cells were reactivated using 1 μg/mL Dox for 24 hours prior to lysis and immunoblotting using antisera that recognizes DR1 in both KapB and KapC. Specific antibody to β-actin served as a loading control. One representative experiment of three is shown. **(B-C)** BAC16-iSLK cells were reactivated with 1 μg/mL Dox and 1 mM NaB for 0, 24, 48, 72, or 96 hours. At each time point cells were trypsinized, pelleted and lysed to extract DNA. qPCR was performed using ORF26 and host β-actin specific primers. Values represent either fold-change relative to time 0 (latency) for each recombinant virus (B) *n*=3, mean ± SD, or fold-change relative to WT at 0, 24, 48, 72 and 96h post reactivation, (C) *n*=3; mean ± SD, (*, P < 0.05; **, P < 0.01; ***, P < 0.001; ****, P < 0.0001; ns, nonsignificant). **(D).** BAC16-iSLK cells were reactivated as above for 96h, virus-containing supernatant was collected, clarified and DNase-treated prior to extraction of virion-protected DNA. qPCR was performed using ORF26 and luciferase (control) specific primers. Values represented fold-change relative to WT. *n*=3; mean ± SD, (*, P < 0.05; **, P < 0.01; ***, P < 0.001; ****, P < 0.0001; ns, nonsignificant). **(E)** BAC16-iSLK cells were reactivated for 96 hours. Virus-containing supernatant for each recombinant cell line was then collected, clarified, and diluted 1:4 with fresh DMEM and used to infect naïve 293T cells. After 24h, cells were fixed, permeabilized and stained using antisera specific for viral LANA (green) and Hoescht (nucleus, blue). Images were captured using Zeiss AxioObserver Z1 microscope with the 40X objective. Scale bar = 20 μm. One representative experiment is shown.

To determine if each of our kaposin-deficient iSLK cell lines was capable of reactivation, completion of a full lytic replication cycle, and the production of progeny virions, we reactivated the cell lines with Dox, to activate RTA expression, and NaB and measured intracellular genome copy number by qPCR at 24, 48, 72, and 96 hours post reactivation. Genome copy number at each time point was normalized to its matched starting genome copy number in latency since each of the latent cell lines had a different episome copy number (Figure 3F) prior to reactivation. For all recombinants, viral genome copy number significantly increased between 48 and 72 hours after reactivation (Figure 5B). WT and ΔKapB viruses exhibited ∼15-fold increase in intracellular genome copy number over latent copy number, ΔKapABC and ΔKapC viruses demonstrated a ∼40-fold increase, and ΔKapBC virus genome copy number increased ∼200-fold by 96 hours post reactivation. We then determined how the reactivation process altered viral genome copy number for each kaposin-deficient recombinant relative to WT and observed that all kaposin-deficient viruses except for ΔKapB reached the same relative genome copy number as WT by 96 hours post reactivation despite starting with less genomes than WT (Figure 5C). These data show that all kaposin-deficient viruses are capable of genome replication to increase viral DNA copy number after reactivation. Because the starting point of the reactivation process was unequal (WT latent iSLK cells have more viral episomes at the ‘starting point’ of reactivation than any of the kaposin-deficient iSLK cell lines, including ΔKapBC latent iSLK cells, Figure 3G), we were hesitant to draw quantitative conclusions about the efficiency of genome replication after reactivation between our panel of kaposin-deficient viruses.

We next determined if kaposin-deficient reactivated iSLK cells released progeny virus particles into the media. To do so, we harvested the virus-containing supernatant at 96 hours post reactivation and subjected it to DNase digestion to eliminate unencapsidated viral DNA, then extracted the capsid-protected DNA and conducted qPCR for viral DNA (Figure 5D). We found that kaposin-deficient viruses released similar numbers of DNase-protected viral genomes to WT (Figure 5D). Although this observation suggests that the ability to package genomic material, assemble and release progeny virus does not require the *kaposin* locus, we and others have observed that extracellular genome copy number does not always correlate with infectious virus (21). To determine whether released virus from each kaposin-deficient latent iSLK cell line were competent to infect naïve cells, we incubated a naïve monolayer of 293T cells with supernatant from reactivated iSLK cells and 24 hours later, stained cells for the latent protein LANA. We identified LANA-expressing cells in all 293T cell monolayers infected by the panel of kaposin-deficient recombinant viruses, (Figure 5E) showing that each kaposin-deficient virus can produce infectious progeny virions. This is consistent with our earlier observations using the 293T-iSLK co-culture system, as this method of latent cell line creation relies on successful lytic replication and virion production.

### Latent iSLK cells that lack Kaposin B display altered latency phenotypes

Previous work from our lab used ectopic overexpression of KapB to characterize its function independent of viral infection. Consequently, we were most interested in analyzing the behaviour of the ΔΚ;apB recombinant virus. We reactivated WT and ΔKapB-BAC16 iSLKs to produce progeny virus, equalized extracellular genomic copy number, and infected naïve cells for 24 hours; however, we saw very few LANA-positive cells post infection (Figure 6A). Given the inconsistencies between stable BAC16 cell lines reported by others (79, 80), we decided to re-derive another ΔKapB-latent iSLK cell line. Using the same population of ΔKapB-BAC16-transfected 293Ts cells and an identical protocol for co-culture, selection, and expansion, we produced a new ΔKapB iSLK cell line that we will refer to as ΔKapB-2 iSLKs, whereas our original ΔKapB iSLK cell line will be referred to as ΔKapB-1. Following ΔKapB-2 iSLK reactivation, we noticed a greater cytopathic effect in reactivated ΔKapB-2 iSLKs than previously observed in reactivated ΔKapB-1 iSLKs. We then equalized extracellular particle number as above, infected naïve cells with ΔKapB-2 BAC16 and observed a 2-fold reduction in LANA-positive cells compared to WT virus (Figure 6B), a great improvement compared to ΔKapB-1. The large difference between infectious progeny production from ΔKapB-1 versus ΔKapB-2 latent iSLKs prompted us to derive two additional ΔKapB iSLK cell lines as described above, termed ΔKapB-3 and ΔKapB-4. We compared several features between these four ΔKapB iSLK cell lines. We noted that although all four latent cell lines were RFP-positive, ΔKapB-1 latent iSLK cells did not fluoresce as brightly as WT or the other ΔKapB latent cell lines (Figure 6C). We stained these latent cells for LANA nuclear bodies (NBs) and observed several differences between LANA NBs present in latent WT iSLKs compared to the four ΔKapB latent iSLK cell lines (Figure 6D). Although WT latent cells had fewer LANA NBs per cell than all four ΔKapB latent cell lines (Figure 6E), WT LANA NBs averaged a much larger area than in ΔKapB latent cell lines (Figure 6F). Of the four ΔKapB latent cell lines, ΔKapB-4 displayed LANA NB character that most closely resembled WT; namely, fewer LANA NBs that were brighter and larger (Figure 6D-F). However, all four ΔKapB latent cell lines showed episome copy number ∼10-25% that of WT latent iSLKs (Figure 6G). Of these, ΔKapB-2 and ΔKapB-4 had the higher genome copy number, consistent with ΔKapB-2 iSLK improved reactivation (ΔKapB-2, Figure 6B) and ΔKapB-4 iSLK LANA NB character (ΔKapB-4, Figure 6D-F). In addition, all four ΔKapB latent cell lines displayed reduced steady-state levels of LANA protein compared to WT (Figure 6H). Taken together, these distinct differences clustered the four ΔKapB iSLK latent cell lines as defective compared to WT in terms of key latency phenotypes such as LANA NB area, steady-state levels of LANA protein, and KSHV genome copy number.

**Figure 6.**
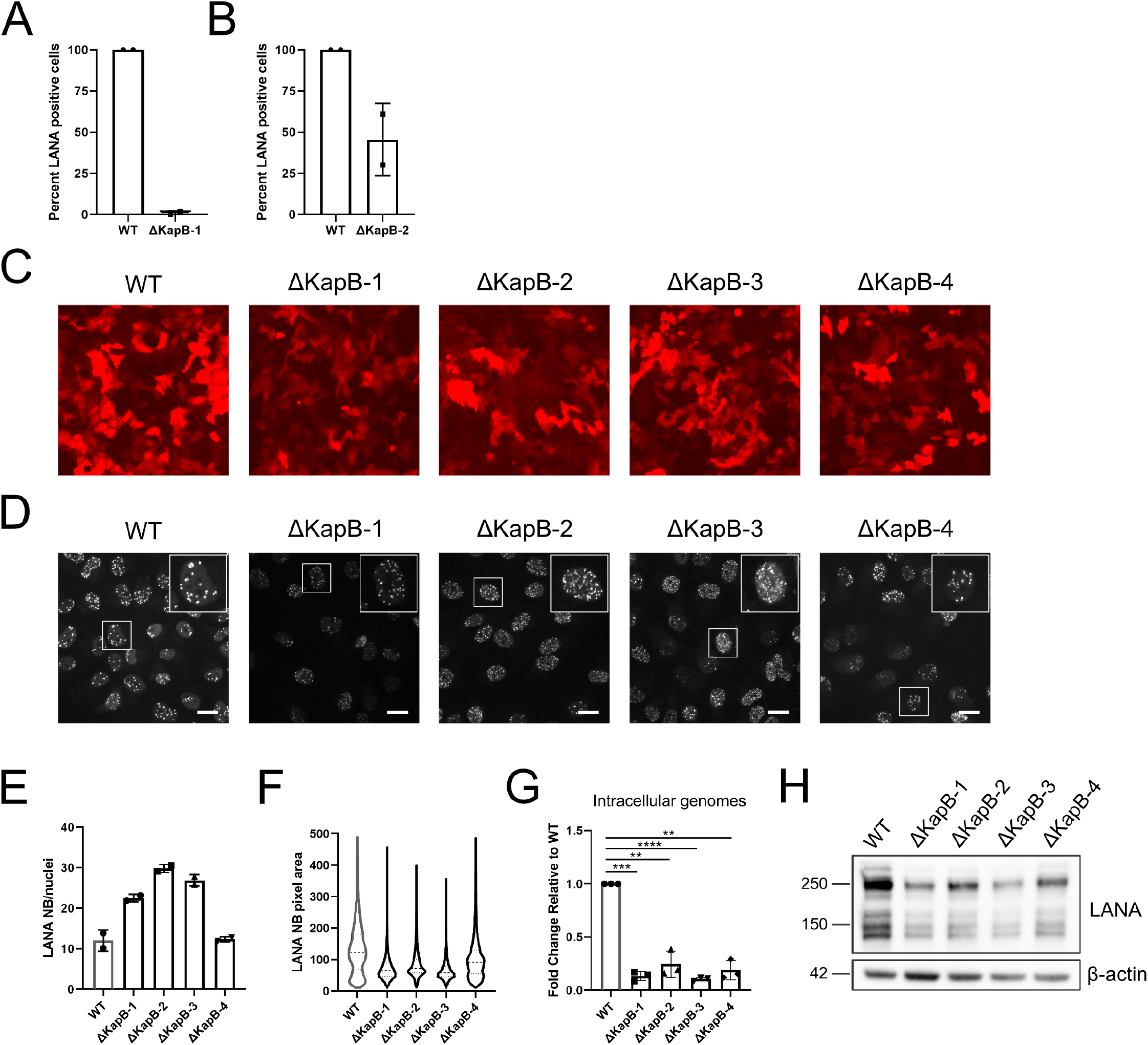
Primary infection of endothelial cells with BAC16ΔKapB virus is impaired. **(A-B)** Equal numbers of virus particles (based on DNase-protected extracellular genome copy number) of WT and either ΔKapB-1 and ΔKapB-2 were used to infect HUVECs. 24h later, cells were fixed, permeabilized and immunstained for viral LANA, then stained with Hoescht. The number of LANA-expressing cells was quantified and is represented as a percent of the total number of cells counted. *n*=2; mean ± SD. **(C)** Images of four different ΔKapB BAC16-iSLK latent cell lines and WT BAC16-iSLKs were captured using an EVOS-FL microscope with the 10X objective. One representative experiment of two is shown. **(D-F)** Latent ΔKapB and WT BAC16-iSLKs were fixed, permeabilized and immunostained for viral LANA, then stained with Hoescht. LANA-NBs were visualized (D), and the number of LANA-NBs per cell was counted using CellProfiler (E) *n*=2; mean ± SD. LANA-NB area was calcuated for at approximently 4000 individual LANA-NBs per cell line and the distribution of area shown in (F). **(G)** Latent ΔKapB and WT BAC16-iSLKs were trypsinized, pelleted and lysed to extract DNA. qPCR was performed using ORF26 and β-actin specific primers. Values are represented as fold change relative WT. *n*=3, mean ± SD, (*, P < 0.05; **, P < 0.01; ***, P < 0.001; ****, P < 0.0001; ns, nonsignificant). **(H)** Latent BAC16-iSLKs were lysed and harvested for immunoblotting using LANA and β-actin specific antibodies. One representative experiment of two is shown.

To determine if we could restore the defects associated with LANA expression in the established ΔKapB iSLK cell lines, we used lentiviruses to ectopically express KapB or an empty vector control (Figure 7A-C). We used RT-qPCR to measure LANA transcript levels with and without complementation. ΔKapB latent iSLK cell displayed ∼10-40% of the LANA RNA transcript levels detected in WT iSLKs (Figure 7A). Overexpression of KapB failed to alter LANA transcript levels compared to the control in any of the ΔKapB iSLK cell lines (Figure 7B). Moreover, latent ΔKapB iSLK cell lines showed reduced steady-state levels of LANA protein, as above, that was again not restored by providing KapB *in trans* (Figure 7C). To determine if complementation with KapB prior to primary infection and latency establishment could restore these defects, we transduced naïve iSLK cells with KapB-expressing or control lentiviruses prior to primary infection with ΔKapB. KapB expression did not complement the low genome copy number or reduced LANA protein level seen in ΔKapB-infected cells after primary infection (Figure 7D-F). To test if KapB could complement primary infection of another cell type, we transfected 293Ts with two different versions of KapB (KS lung and BAC16) prior to primary infection. Neither version of KapB was able to rescue low KSHV genome copy number or LANA protein levels after *de novo* infection of 293Ts (Figure 7G-I). Taken together, these data suggest that an aspect of the kaposin locus is required *in cis*, as providing the KapB polypeptide *in tran*s, via non-targeted lentiviral integration, cannot complement the defects associated with ΔKapB latency after primary infection.

**Figure 7.**
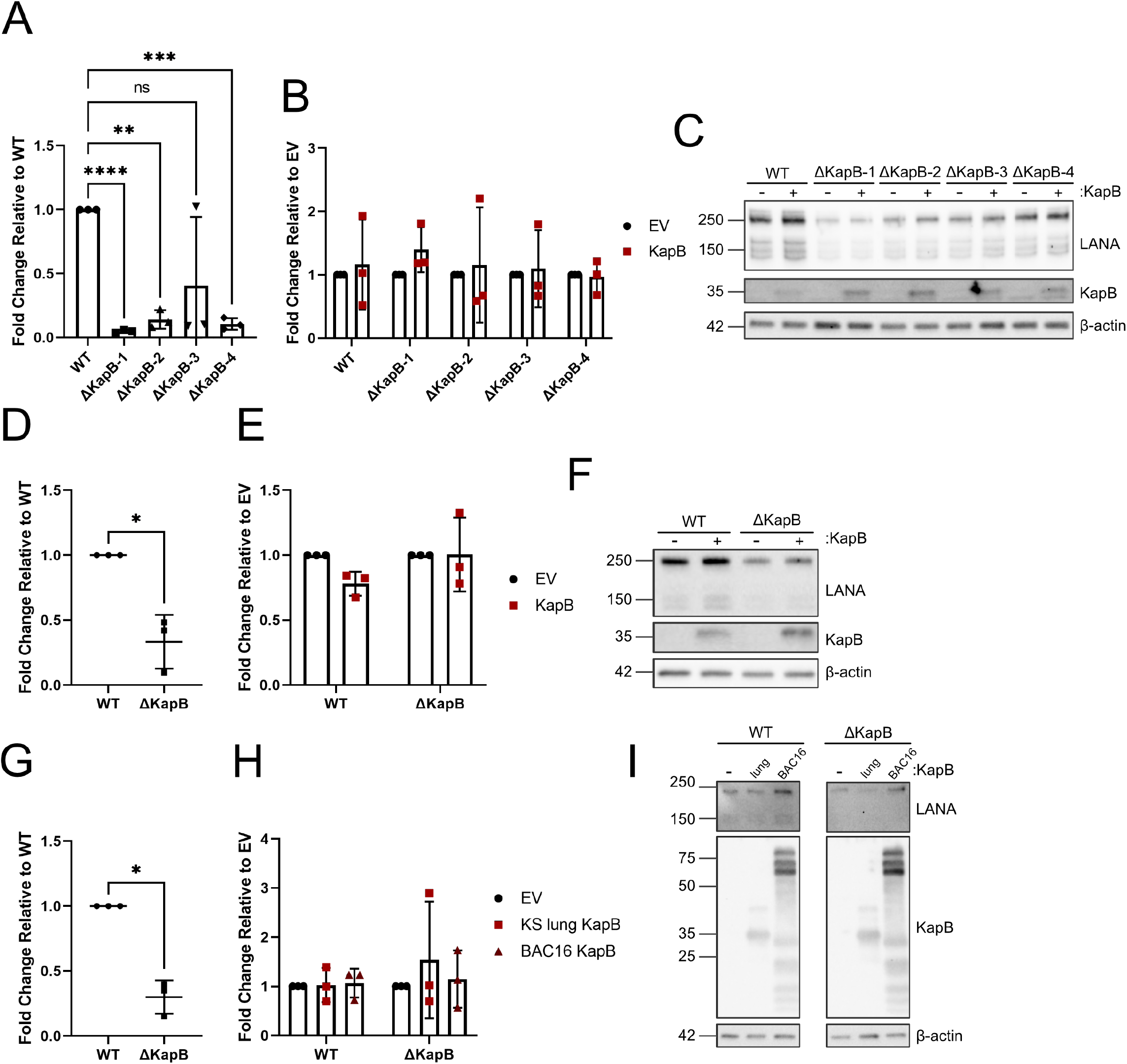
Ectopic expression of KapB does not rescue altered latency phenotypes in ΔKapB infected cells. **(A-C)** Latent WT or ΔKapB BAC16-iSLK cells lines were transduced to express either an empty vector (EV) control or KapB (KS lung isolate). 24h later, cells were selected using blasticidin for 48h prior to seeding. The following day seeded cells were lysed and harvested for RNA or protein analysis. (A-B) RT-qPCR was performed using LANA and 18S specific primers. *n*=3; mean ± SD, (*, P < 0.05; **, P < 0.01; ***, P < 0.001; ****, P < 0.0001; ns, nonsignificant). In A, LANA RNA transcript levels in ΔKapB iSLK cell lines are shown normalized to WT iSLK cells. In B, LANA RNA transcript levels are shown after KapB complementation, normalized to each respective EV control. (C) Immunoblotting was performed using LANA, DR1 (KapB) and β-actin specific antibodies. One representative experiment of three is shown. **(D-F)** Naïve iSLK cells were transduced with EV or KapB lentiviruses as in (A). After seeding, cells were infected with equal particle numbers of WT and ΔKapB BAC16 viruses as determined via DNase-protected extracellular genome copy number. 24h post infection, cells were either trypsinized, pelleted and lysed to extract DNA or lysed for immunoblotting. (D-E) qPCR was performed for KSHV DNA using ORF26 and host β-actin specific primers. *n*=3; mean ± SD, (*, P < 0.05; **, P < 0.01; ***, P < 0.001; ****, P < 0.0001; ns, nonsignificant). In D, Viral genome copy number for iSLKs after *de novo* infection with ΔKapB is normalized to the WT control. In E, viral genome copy number for each virus infection after complementing with KapB is normalized to each respective EV control. (F) Immunoblotting was performed using LANA, DR1 (KapB) and β-actin specific antibodies. One representative experiment of three is shown. **(G-I)** 293T cells were transfected with three different ectopic expression constructs for KapB: KapB derived from KS lung (45) and KapB BAC16. 24 h after transfection, cells were infected with equal particle numbers of WT and ΔKapB BAC16 viruses and processed as above for qPCR (G-H, *n*=3; mean ± SD, (*, P < 0.05; **, P < 0.01; ***, P < 0.001; ****, P < 0.0001; ns, nonsignificant). or immunoblotting (I, n=1). In G, Viral genome copy number for 293Ts after *de novo* infection with ΔKapB is normalized to the WT control. In H, viral genome copy number for each virus infection after complementing with different versions of KapB is normalized to each respective EV control.

### KapB is required for processing body disassembly after *de novo* infection of HUVECs

We have previously shown that ectopic expression of KapB in human umbilical vein endothelial cells (HUVECs) is sufficient to recapitulate key phenotypes associated with KSHV latency, including the disassembly of processing bodies (PBs), cytoplasmic RNA and protein granules whose loss correlates with enhanced inflammatory cytokine RNAs (51, 53–57). To determine if KapB was necessary for KSHV-induced PB loss, we reactivated ΔKapB-2 latent iSLK cells and used this virus to infect HUVECs. After 96 hours post infection, we fixed the infected cells and stained the monolayer for both LANA and for a PB-resident protein called Hedls/EDC4 (81, 82). We observed that ∼80-90% of the cell monolayer was infected with either ΔKapB BAC16 or WT BAC16 virus. In LANA-positive cells that were infected with WT virus, we observed a two-fold decrease in PBs per infected cell compared to the mock-infected control (Figure 8A-B), as shown previously (51). In LANA-positive cells infected with ΔKapB-2 virus, PB numbers were not significantly altered relative to the mock-infected control (Figure 8A-B). We also confirmed that WT-infected HUVECs expressed KapB, but that KapB was absent in ΔKapB-infected HUVECs (Figure 8C). These data reveal for the first time that the KapB protein is not only sufficient but also necessary for KSHV-induced PB disassembly following *de novo* infection.

**Figure 8.**
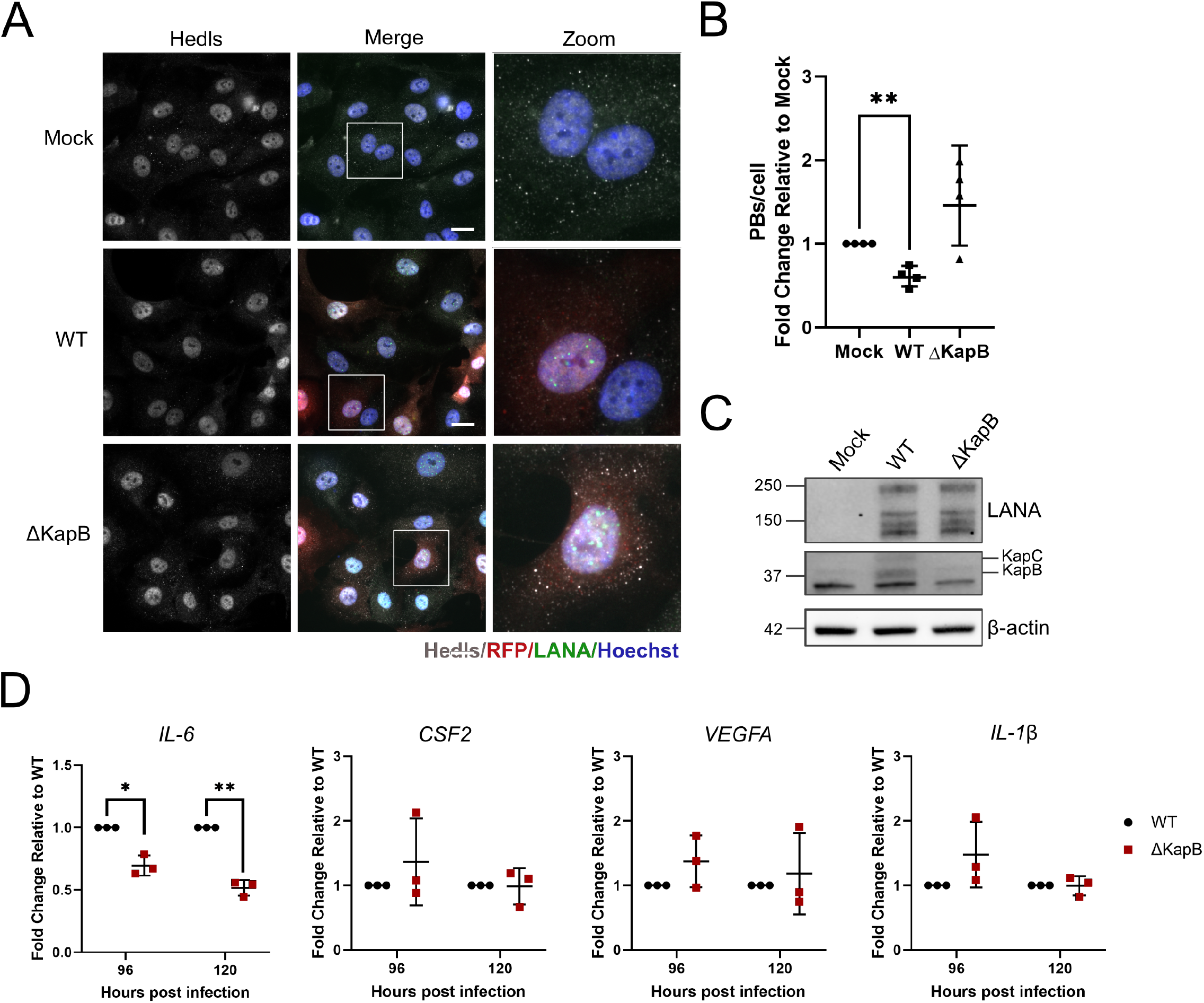
KapB is required for PB disassembly following *de novo* HUVEC infection. **(A)** WT and ΔKapB BAC16 viruses were titrated using 293T cells and RFP expression was used to estimate an equal number of infectious virions for subsequent *de novo* infection of HUVECs. At 96hpi, HUVECs were fixed, permeabilized and immunostained for either viral LANA (green), the PB-resident protein Hedls/EDC4 (white) then stained with Hoescht (nucleus, blue). Images were captured using Zeiss AxioObserver Z1 microscope with the 40X objective. Scale bar = 20 μm. **(B)** Hedls puncta per cell were quantified using CellProfiler. For mock infections, all cells were counted; for WT and ΔKapB-infected cells, only LANA-positive cells were counted. Data is represented as fold change in PBs per cell relative to mock infection. *n*=4; mean ± SD, (*, P < 0.05; **, P < 0.01; ***, P < 0.001; ****, P < 0.0001; ns, nonsignificant). **(C)** Cells infected as in (A) were lysed at 96 hpi and harvested for immunoblotting using LANA, DR1 (KapB and C) and β-actin specific antibodies. One representative experiment of three is shown. **(D)** Cells infected as in (A) were lysed at 96 and 120 hpi and harvested for RNA extraction and RT-qPCR using IL-6, CSF2, VEGFA, IL-1β and HPRT specific primers. *n*=3; mean ± SD, (*, P < 0.05; **, P < 0.01; ***, P < 0.001; ****, P < 0.0001; ns, nonsignificant).

PBs constitutively repress or degrade cytokine transcripts that contain AU-rich elements (AREs); accordingly, we and others have observed that PB disassembly correlates with enhanced steady-state levels of some cytokine transcripts (51, 53–57). After primary infection of HUVECs with WT or ΔKapB, we measured steady-state levels of selected ARE-containing cytokine transcripts by RT-qPCR (Figure 8D). We predicted that if PBs remain intact, as is after ΔKapB infection, cytokine levels would be decreased relative WT-infected cells as the ARE-RNAs from which they are expressed would be subject to PB-mediated decay. Consistent with this hypothesis, after infection with ΔKapB, IL-6 transcript levels were much reduced relative to that observed with WT infection (Figure 8D). This is the expected response for a cytokine mRNA that is shuttled to PBs for constitutive decay. However, we did not observe a strong difference between steady-state levels of other ARE-containing cytokine transcripts (VEGF, IL-1β, GM-CSF) after WT versus ΔKapB infection (Figure 8D), as not all ARE-RNAs respond in the same manner to altered PB dynamics, as shown in (83). Moreover, IL-6 mRNA levels are extremely sensitive to changes in PBs (84) and we previously observed IL-6 transcript levels increase in response PB disassembly induced by KapB (85). Taken together, these data show that KSHV-induced PB disassembly correlates with ∼2-3-fold increased steady-state level of the IL-6 transcript and that both PB disassembly and enhanced IL-6 RNA levels are absent in cells infected with ΔKapB.

We previously showed that ectopic expression of KapB is sufficient to induce primary ECs to elongate and spindle, thereby recapitulating the morphology of infected cells within KS lesions (51, 86). However, others have observed that ectopic expression of v-FLIP likewise elicits EC spindling (87) and that the deletion of v-FLIP from the viral genome eliminates EC spindling, although this effect is overcome at high multiplicity of infection (88). To test if KapB is necessary for EC spindling, we used WT of ΔKapB BAC16, derived as above, to infect HUVECs, fixed cells at 96hpi and visualized spindling at low magnification by brightfield microscopy or at high magnification by staining fixed cells with phalloidin to label actin filaments. Robust cell spindling was observed in LANA-positive, WT-infected cells (Figure 9). We also observed spindled cells in LANA-positive, ΔKapB-infected HUVECs (Figure 9). We conclude that although ectopic expression of KapB is sufficient to elicit EC spindling (51), it is not necessary for EC spindling during KSHV latency.

**Figure 9.**
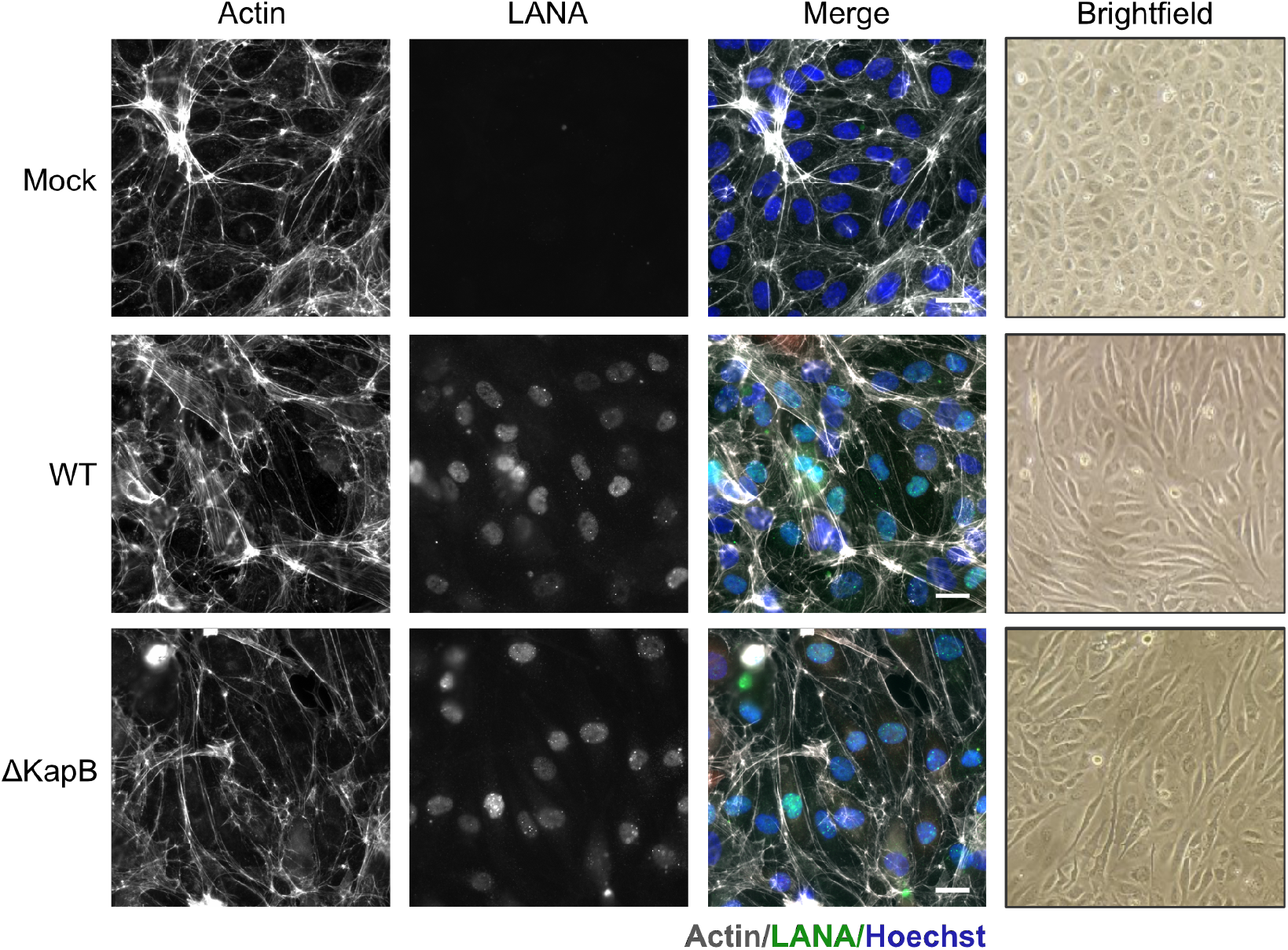
KapB is not required for endothelial cell spindling following de novo HUVEC infection. WT and ΔKapB BAC16 viruses were titrated using 293T cells and RFP expression was used to estimate an equal number of infectious virions for subsequent *de novo* infection of HUVECs, as in Figure 8. At 96 hpi, brightfield images were acquired at 10X, cells were fixed, permeabilized and immunostained for viral LANA (green). Phalloidin and Hoescht staining was used to visualize actin (white) and nuclei (blue), respectively. Images were captured using Zeiss AxioObserver Z1 microscope with the 40X objective. Scale bar = 20 μm. One representative image of three experiments is shown.

## Discussion

The kaposin locus comprises a significant portion of the coding capacity of the KSHV latency locus, yet its role in KSHV latent and lytic replication is not fully appreciated. Prior to this report, the KSHV field lacked recombinant viruses to enable functional analysis of individual kaposin proteins in the context of KSHV infection. Here we described the construction and validation of a suite of kaposin-deficient recombinant viruses using the BAC16 bacterial artificial chromosome and a panel of corresponding stable iSLK cell lines. We deleted or recoded kaposin ORFs individually or in combination, generating several mutants: ΔKapABC, ΔKapBC, ΔKapB, and ΔKapC. We showed that all kaposin-deficient viruses established latency in iSLKs, replicated the viral genome upon reactivation, and produced progeny virions that could be used for *de novo* infection of naïve cells. Despite this, all kaposin-deficient latent iSLK cells lines displayed reduced viral genome copy number during latency compared to WT. In addition, we characterized four derivations of ΔKapB-iSLKs and showed that these cells not only had markedly reduced genome copy number, but also reduced LANA NB area and LANA RNA and protein levels. Attempts to complement the reduced expression of LANA in ΔKapB-iSLKs by ectopic expression of KapB had no effect, nor did attempts to complement the reduced KSHV genome copy number after primary infection with ΔKapB virus. Together, these data suggest that the kaposin locus plays a role in latency establishment that cannot be restored by complementing with KapB protein *in trans*. Finally, we used ΔKapB BAC16 to show that KapB is not only sufficient but also necessary for KSHV-mediated PB disassembly following *de novo* infection of primary ECs, verifying the contribution of an individual kaposin protein to KSHV biology. This toolkit will allow us to understand the role of the kaposin locus in KSHV replication with unprecedented precision, making it a significant advance for the field.

Previous knowledge about the role of the kaposin locus was derived from studies using the ectopic expression of individual kaposin ORFs (47, 51, 52, 89–91). To date, no comprehensive kaposin-deficient BAC16 studies have been published. Toth *et al.*, created a kaposin knockout BAC16 virus by deleting a 1319-nucleotide region that included the coding region of KapA and the majority of the coding regions of KapB and KapC (92). This recombinant virus was used to test for the recruitment of proteins from polycomb repressive complexes 1 and 2 to two lytic viral promoters after primary infection of SLK cells (92). For this phenotype, the kaposin knockout virus displayed no difference from WT, but aside from this, no additional analyses related to this mutant were reported. Gallo *et al.* created a K12 mutant in which a kanamycin cassette disrupts the K12 ORF, but reported limited phenotypic analysis related to this virus (93). Finally, Campbell *et al.* investigated the role of the RTA-responsive element (RRE) promoter in the kaposin locus, and in so doing created a K12 RRE-mutant; however, this study focused solely on RTA-recruitment and genomic architecture (94). In the current study, our recombinant viruses were created by removing all DR1 and DR2 sequences (ΔKapBC) with or without mutagenesis of the KapA ORF start codon (ΔKapABC), or by replacing the DR region with alternatively coded nucleotide sequences (ΔKapB and ΔKapC). Our recoding strategy had several advantages. First, the translation potential of the *kaposin* locus was reduced by insertion of multiple termination codons, eliminating expression of some protein products while maintaining other ORFs by alternative codon usage. Second, recoding destroyed the repetitive nature of the locus and decreased the GC-content, making the region much more amenable to error-free homologous recombination. Third, in all of the recombinant kaposin-mutant BACs, regulatory sequences for transcription and miRNA processing were left intact. We also created corresponding latent iSLK cell lines for the kaposin-deficient BAC16 constructs. All four of these kaposin-deficient latent iSLK cell lines showed drastic reductions in viral genome copy number compared to WT. The cell lines were created by co-culturing BAC16-transfected HEK 293Ts with naïve iSLKs, a widely used method reported to be more efficient than direct transfection of recombinant BAC16 DNA, that leverages cell-to-cell spread of virions from reactivated 293T cells to iSLKs (5, 59, 69, 95). Despite using this strategy, we failed to produce latent cell lines that contained high genome copy numbers for any of the kaposin-deficient BAC16 DNA. We repeated the co-culture procedure for the most impaired of these four times, and each attempt yielded similar results; all four ΔKapB iSLK cell lines displayed reduced viral episome number and altered LANA NBs compared to WT BAC16 cells. Although these data are consistent with the idea that kaposin-deficient viruses are impaired in one or more key steps required for the establishment of latency, further study is required to characterize the nature of this defect.

LANA expression and function is central to the establishment of KSHV latency. LANA forms homo-oligomers and binds the terminal repeat (TR) regions of the KSHV genome while simultaneously binding histones H2A and H2B of host chromatin, thereby tethering the viral DNA to host chromosomes (8, 10, 11, 96). This binding leads to the formation of structures called LANA NBs that colocalize with viral genomes (10, 97, 98). The number of LANA-positive puncta in an infected cell correlates with the number of viral episomes (13). KSHV *de novo* infection of a cell monolayer is heterogenous and results in variable levels of intracellular viral genomes between individual latent cells, and this heterogeneity is also represented by diverse LANA NB number per cell (13). Therefore, robust LANA NB staining is a marker for successful latency establishment. We and others have observed heterogenous LANA NB size within and between latent infected cells; however, the significance of this size variability is less clear. Because LANA can bind to nucleosomes in cellular chromosomes as well as those bound to viral episomes (15), LANA NBs are classified as supramolecular structures that cluster multiple viral episomes together (12, 98). Genome clustering was proposed to be an effective evolutionary strategy that allows KSHV to rapidly increase viral genome copy number per cell at the expense of total number of cells infected (15). It is not yet clear if a minimum number of viral episomes are required to create a supramolecular LANA NB after *de novo* infection; however, WT KSHV does preferentially form clustered LANA NBs (15). We have observed that latent ΔKapB iSLK cells possess smaller and more uniform LANA NBs compared to those found in WT iSLK cells. Latent ΔKapB iSLK cells express lower amounts of LANA RNA and protein, a phenotype that we could not complement by ectopic expression of KapB *in trans*. Therefore, we report an additional correlative phenotype between low genome copy number, lower LANA expression and smaller LANA NBs observed in multiple derivations of ΔKapB latent iSLK cell lines and speculate that ΔKapB iSLK cells do not contain enough viral episomes and/or LANA protein to form large LANA NBs with clustered genomes.

The kaposin locus overlaps with one of two possible viral origins of lytic genome replication, called OriLytR. Though very few studies have investigated the contribution of OriLytR/kaposin to viral genome replication, detailed analyses on the function of OriLytL have revealed that it facilitates genome copying after viral reactivation and upon *de novo* infection (99–102). More recent work has shown an essential role for OriLytL transcription in viral genome replication after *de novo* infection (102). We speculate that loss of the GC-rich kaposin repeat region in our mutants impairs the contribution of OriLytR to viral genome copying that occurs after *de novo* infection, as was observed for a recombinant virus lacking OriLytL (102). A reduction in viral genome copy number after *de novo* infection would be predicted to impair latency establishment and diminish the clustering of viral episomes into larger LANA NBs, which may explain the different LANA NB phenotype we observed in all four ΔKapB iSLK latent cell lines. Since we failed to complement the low genome copy number observed after primary infection with ΔKapB, these data suggest that the role of the *kaposin* locus after primary infection is derived from either the DNA locus or its RNA transcript *in cis* and not from the polypeptide provided *in trans*. At odds with our model is that the loss of the kaposin region does not appear to negatively impact viral genome replication after reactivation. We speculate that this step is not as sensitive to the loss of one of two functional origins of lytic replication, as also observed for loss of OriLytL (120). Further experimentation will be needed to reveal the precise mechanism by which kaposin region may promote genome amplification after primary infection and how this alters LANA NB formation, function, and latency establishment.

KSHV latency predominates in most cells in culture and in KS tumours, suggesting viral proteins expressed at this time are important in promoting tumourigenesis. We have previously shown that ectopic expression of KapB in primary ECs recapitulates two features of the KS lesion: cell spindling and elevated inflammatory cytokine expression via PB disassembly. Using the ΔKapB BAC16 recombinant virus, we analyzed if KapB was not only sufficient but necessary for these KSHV-associated tumor phenotypes. We show that after *de novo* infection of HUVECs, ΔKapB BAC16 cells can induce cell spindling to a similar extent as WT BAC16 virus. These data are consistent with reports that the viral latent gene vFLIP is also capable of causing endothelial cell spindling; therefore, the loss of KapB does not eliminate the spindling phenotype (87). We propose that KapB and v-FLIP may act synergistically to promote EC spindling during KSHV latency, and that in absence of one, the other is able to compensate. We also used the ΔKapB BAC16 recombinant virus to determine if KapB was necessary for KSHV-induced PB disassembly. PBs are cytoplasmic sites for the constitutive turnover or translational suppression of ARE-containing inflammatory cytokine mRNAs (51, 53–57). We previously showed that latent KSHV infection of primary ECs caused PB disassembly (51). Here, we also observed that 96 hours after d*e novo* infection of HUVECs with WT BAC16, PBs were disassembled. However, d*e novo* infection of HUVECs with ΔKapB BAC16 did not cause PB disassembly, and steady-state levels of IL-6 RNA, an ARE-containing cytokine transcript known to be regulated by PB levels, were reduced relative to that after WT infection (84). This reveals that KapB is both necessary and sufficient for PB disassembly during KSHV latency and represents the first time that a phenotypic function can be specifically attributed to a single kaposin protein in the context of KSHV infection.

Our group and others showed that PBs also decrease during lytic replication and that more than one viral protein is sufficient for this effect (53, 103). Here, we observed PB disassembly 96 hours after *de novo* infection of HUVECs, and we have previously shown PB disassembly in latent HUVECs (51). In contrast, Sharma *et al.* reported that PBs were not lost in a KSHV-infected latent primary effusion lymphoma cell line. This discrepancy could be attributed to the use of B-lymphocytes in their studies or their visualization of PBs by staining largely for the PB-resident protein GW182, an antigen that has been shown to adopt a punctate pattern distinct from other PB marker proteins (103, 104). Taken together, the observation that KSHV encodes multiple proteins in both phases of its replication to elicit PB disassembly suggests the functional relevance of this phenotype for viral infection. Because PBs constitutively repress many pro-inflammatory, angiogenic, or pro-tumourigenic cellular transcripts, we propose that the absence of microscopically visible PBs may contribute to the inflammatory and angiogenic KS microenvironment and that KapB is necessary for this effect in KS lesions.

In summary, this work describes the construction and preliminary analysis of four different kaposin-deficient BAC16 viruses and corresponding iSLK latent cell lines. We used these mutants to specifically attribute the phenotype of PB disassembly following *de novo* infection of primary to the viral protein KapB. This work also provides preliminary data to suggest that the *kaposin* locus aids in the establishment of KSHV latency after primary infection, a role that may suggest that the region derives function from its GC-rich direct repeats in addition to its protein products. This is exciting because it suggests that the *kaposin* locus is truly multipurpose, deriving distinct functions important for viral replication from its RNA transcript, and its protein products. We hope these mutants will enable future studies to better evaluate the roles of individual kaposin protein products during infection. Moreover, we believe our strategy to recode protein products in multiple reading frames can serve as a mutagenesis roadmap for understanding other complex, polycistronic viral loci.

## Acknowledgements

We sincerely thank the members of the Corcoran lab for helpful discussions about this work including Ms. Julie Ryu for designing recoded ΔKapB sequence. We would like to thank Dr. Craig McCormick (Dalhousie University) for valuable feedback on this manuscript and the McCormick lab for plasmids, expertise and advice. We would like to thank Dr. Anne Vaahtokari of the Charbonneau Microscopy Facility, UCalgary for microscopy support. We would like to thank Dr. John Archibald (Dalhousie University) and the Archibald lab for help performing pulse-field gel electrophoresis and Dr. Jae Jung (Cleveland Clinic) for sharing the BAC16-RFP bacterial artificial chromosome. MK was supported by a CSM graduate training award, a CIHR CGS-M scholarship, and a CIHR doctoral award. GM was supported by a CRTP graduate training award. Operating funds to support this work derive from a CIHR Project Grant PJT-153210 to JAC.

## Author Contributions

Mariel Kleer: Conceptualization, Experimentation, Analysis, Paper Writing

Grant MacNeil: Conceptualization, Experimentation, Analysis

Eric S. Pringle: Conceptualization, Supervision

Jennifer A. Corcoran: Conceptualization, Experimentation, Supervision, Funding Acquisition, Project Administration, Paper Writing

## References

1. Chang Y, Cesarman E, Pessin MS, Lee F, Culpepper J, Knowles DM, Moore PS. 1994. Identification of herpesvirus-like DNA sequences in AIDS-associated Kaposi’s sarcoma. Science (80-) https://doi.org/10.1126/science.7997879.

2. Cesarman E, Chang Y, Moore PS, Said JW, Knowles DM. 1995. Kaposi’s sarcoma— associated herpesvirus-like DNA sequences in AIDS-related body-cavity—based lymphomas. N Engl J Med https://doi.org/10.1056/NEJM199505043321802.

3. Soulier J, Grollet L, Oksenhendler E, Cacoub P, Cazals-Hatem D, Babinet P, D’Agay MF, Clauvel JP, Raphael M, Degos L, Sigaux F. 1995. Kaposi’s sarcoma-associated herpesvirus-like DNA sequences in multicentric Castleman’s disease. Blood https://doi.org/10.1182/blood.v86.4.1276.bloodjournal8641276.

4. Lagunoff M, Bechtel J, Venetsanakos E, Roy AM, Abbey N, Herndier B, Ganem D, McMahon M, Ganem D. 2002. De Novo Infection and Serial Transmission of Kaposi’s Sarcoma-Associated Herpesvirus in Cultured Endothelial Cells. J Virol https://doi.org/10.1128/JVI.76.5.2440.

5. Renne R, Blackbourn D, Whitby D, Levy J, Ganem D. 1998. Limited transmission of Kaposi’s sarcoma-associated herpesvirus in cultured cells. J Virol https://doi.org/10.1128/JVI.72.6.5182-5188.1998.

6. Purushothaman P, Thakker S, Verma SC. 2015. Transcriptome Analysis of Kaposi’s Sarcoma-Associated Herpesvirus during De Novo Primary Infection of Human B and Endothelial Cells. J Virol https://doi.org/10.1128/jvi.02507-14.

7. Krishnan HH, Naranatt PP, Smith MS, Zeng L, Bloomer C, Chandran B. 2004. Concurrent Expression of Latent and a Limited Number of Lytic Genes with Immune Modulation and Antiapoptotic Function by Kaposi’s Sarcoma-Associated Herpesvirus Early during Infection of Primary Endothelial and Fibroblast Cells and Subsequent Decline of L. J Virol https://doi.org/10.1128/jvi.78.7.3601-3620.2004.

8. Cotter MA, Robertson ES. 1999. The latency-associated nuclear antigen tethers the Kaposi’s sarcoma-associated herpesvirus genome to host chromosomes in body cavity-based lymphoma cells. Virology https://doi.org/10.1006/viro.1999.9999.

9. Szekely L, Kiss C, Mattsson K, Kashuba E, Pokrovskaja K, Juhasz A, Holmvall P, Klein G. 1999. Human herpesvirus-8-encoded LNA-1 accumulates in heterochromatin-associated nuclear bodies. J Gen Virol https://doi.org/10.1099/0022-1317-80-11-2889.

10. Ballestas ME, Chatis PA, Kaye KM. 1999. Efficient persistence of extrachromosomal KSHV DNA mediated by latency-associated nuclear antigen. Science (80-) https://doi.org/10.1126/science.284.5414.641.

11. Schwam DR, Luciano RL, Mahajan SS, Wong L, Wilson AC. 2000. Carboxy Terminus of Human Herpesvirus 8 Latency-Associated Nuclear Antigen Mediates Dimerization, Transcriptional Repression, and Targeting to Nuclear Bodies. J Virol https://doi.org/10.1128/jvi.74.18.8532-8540.2000.

12. De Leo A, Deng Z, Vladimirova O, Chen HS, Dheekollu J, Calderon A, Myers KA, Hayden J, Keeney F, Kaufer BB, Yuan Y, Robertson E, Lieberman PM. 2019. LANA oligomeric architecture is essential for KSHV nuclear body formation and viral genome maintenance during latency. PLoS Pathog https://doi.org/10.1371/journal.ppat.1007489.

13. Adang LA, Parsons CH, Kedes DH. 2006. Asynchronous Progression through the Lytic Cascade and Variations in Intracellular Viral Loads Revealed by High-Throughput Single-Cell Analysis of Kaposi’s Sarcoma-Associated Herpesvirus Infection. J Virol https://doi.org/10.1128/jvi.01156-06.

14. Grundhoff A, Ganem D. 2004. Inefficient establishment of KSHV latency suggests an additional role for continued lytic replication in Kaposi sarcoma pathogenesis. J Clin Invest https://doi.org/10.1172/JCI200417803.

15. Chiu YF, Sugden AU, Fox K, Hayes M, Sugden B. 2017. Kaposi’s sarcoma-associated herpesvirus stably clusters its genomes across generations to maintain itself extrachromosomally. J Cell Biol https://doi.org/10.1083/jcb.201702013.

16. Lukac DM, Renne R, Kirshner JR, Ganem D. 1998. Reactivation of Kaposi’s sarcoma-associated herpesvirus infection from latency by expression of the ORF 50 transactivator, a homolog of the EBV R protein. Virology https://doi.org/10.1006/viro.1998.9486.

17. Lukac DM, Kirshner JR, Ganem D. 1999. Transcriptional Activation by the Product of Open Reading Frame 50 of Kaposi’s Sarcoma-Associated Herpesvirus Is Required for Lytic Viral Reactivation in B Cells. J Virol https://doi.org/10.1128/jvi.73.11.9348-9361.1999.

18. Xu Y, AuCoin DP, Huete AR, Cei SA, Hanson LJ, Pari GS. 2005. A Kaposi’s Sarcoma-Associated Herpesvirus/Human Herpesvirus 8 ORF50 Deletion Mutant Is Defective for Reactivation of Latent Virus and DNA Replication. J Virol https://doi.org/10.1128/jvi.79.6.3479-3487.2005.

19. Sun R, Lin S-F, Staskus K, Gradoville L, Grogan E, Haase A, Miller G. 1999. Kinetics of Kaposi’s Sarcoma-Associated Herpesvirus Gene Expression. J Virol https://doi.org/10.1128/jvi.73.3.2232-2242.1999.

20. Sun R, Lin SF, Gradoville L, Yuan Y, Zhu F, Miller G. 1998. A viral gene that activates lytic cycle expression of Kaposi’s sarcoma-associated herpesvirus. Proc Natl Acad Sci U S A https://doi.org/10.1073/pnas.95.18.10866.

21. Nakajima K, Guevara-Plunkett S, Chuang F, Wang K-H, Lyu Y, Kumar A, Luxardi G, Izumiya C, Soulika A, Campbell M, Izumiya Y. 2020. Rainbow Kaposi’s Sarcoma-Associated Herpesvirus Revealed Heterogenic Replication with Dynamic Gene Expression. J Virol https://doi.org/10.1128/jvi.01565-19.

22. Chen CP, Lyu Y, Chuang F, Nakano K, Izumiya C, Jin D, Campbell M, Izumiya Y. 2017. Kaposi’s Sarcoma-Associated Herpesvirus Hijacks RNA Polymerase II To Create a Viral Transcriptional Factory. J Virol https://doi.org/10.1128/jvi.02491-16.

23. Okoid LM, Kimballid AK, Kasparid RE, Knoxid AN, Coleman CB, Rochford R, Chang T, Alderete B, van Dyk LF, Clambey ET. 2019. Multidimensional analysis of gammaherpesvirus RNA expression reveals unexpected heterogeneity of gene expression. PLoS Pathog https://doi.org/10.1371/journal.ppat.1007849.

24. Zhong W, Wang H, Herndier B, Ganem D. 1996. Restricted expression of Kaposi sarcoma-associated herpesvirus (human herpesvirus 8) genes in Kaposi sarcoma. Proc Natl Acad Sci U S A https://doi.org/10.1073/pnas.93.13.6641.

25. Katano H, Sato Y, Kurata T, Mori S, Sata T. 2000. Expression and localization of human herpesvirus 8-encoded proteins in primary effusion lymphoma, Kaposi’s sarcoma, and multicentric Castleman’s disease. Virology https://doi.org/10.1006/viro.2000.0196.

26. Katano H, Sato Y, Itoh H, Sata T. 2001. Expression of human herpesvirus 8 (HHV-8)-encoded immediate early protein, open reading frame 50, in HHV-8-associated diseases. J Hum Virol.

27. Staskus KA, Gebhard K, Haase AT, Zhong W, Wang H, Renne R, Ganem D, Herndier B, Beneke J, Pudney J, Anderson DJ. 1997. Kaposi’s sarcoma-associated herpesvirus gene expression in endothelial (spindle) tumor cells. J Virol.

28. Bais C, Santomasso B, Coso O, Arvanitakis L, Raaka EG, Gutkind JS, Asch AS, Cesarman E, Gerhengorn MC, Mesri EA. 1998. G-protein-coupled receptor of Kaposi’s sarcoma-associated herpesvirus is a viral oncogene and angiogenesis activator. Nature https://doi.org/10.1038/34193.

29. Montaner S, Sodhi A, Molinolo A, Bugge TH, Sawai ET, He Y, Li Y, Ray PE, Gutkind JS. 2003. Endothelial infection with KSHV genes in vivo reveals that vGPCR initiates Kaposi’s sarcomagenesis and can promote the tumorigenic potential of viral latent genes. Cancer Cell https://doi.org/10.1016/S1535-6108(02)00237-4.

30. Mazzi R, Parisi SG, Sarmati L, Uccella I, Nicastri E, Carolo G, Gatti F, Concia E, Andreoni M. 2001. Efficacy of cidofovir on human herpesvirus 8 viraemia and Kaposi’s sarcoma progression in two patients with AIDS [1]. AIDS.

31. Mesri EA, Feitelson MA, Munger K. 2014. Human viral oncogenesis: A cancer hallmarks analysis. Cell Host Microbe.

32. Martin DF, Kuppermann BD, Wolitz RA, Palestine AG, Li H, Robinson CA. 1999. Oral Ganciclovir for Patients with Cytomegalovirus Retinitis Treated with a Ganciclovir Implant. N Engl J Med https://doi.org/10.1056/nejm199904083401402.

33. Bravo Cruz AG, Damania B. 2019. In Vivo Models of Oncoproteins Encoded by Kaposi’s Sarcoma-Associated Herpesvirus. J Virol https://doi.org/10.1128/jvi.01053-18.

34. Friborg J, Kong WP, Hottlger MO, Nabel GJ. 1999. p53 Inhibition by the LANA protein of KSHV protects against cell death. Nature https://doi.org/10.1038/47266.

35. Fujimuro M, Wu FY, Aprhys C, Kajumbula H, Young DB, Hayward GS, Hayward SD. 2003. A novel viral mechanism for dysregulation of β-catenin in Kaposi’s sarcoma-associated herpesvirus latency. Nat Med https://doi.org/10.1038/nm829.

36. Radkov SA, Kellam P, Boshoff C. 2000. The latent nuclear antigen of Kaposi sarcoma-associated herpesvirus targets the retinoblastoma-E2F pathway and with the oncogene Hras transforms primary rat cells. Nat Med https://doi.org/10.1038/80459.

37. Sin SH, Dittmer DP. 2013. Viral latency locus augments B-cell response in vivo to induce chronic marginal zone enlargement, plasma cell hyperplasia, and lymphoma. Blood https://doi.org/10.1182/blood-2012-03-415620.

38. Qin J, Li W, Gao SJ, Lu C. 2017. KSHV microRNAs: Tricks of the Devil. Trends Microbiol.

39. Arias C, Weisburd B, Stern-Ginossar N, Mercier A, Madrid AS, Bellare P, Holdorf M, Weissman JS, Ganem D. 2014. KSHV 2.0: A Comprehensive Annotation of the Kaposi’s Sarcoma-Associated Herpesvirus Genome Using Next-Generation Sequencing Reveals Novel Genomic and Functional Features. PLoS Pathog https://doi.org/10.1371/journal.ppat.1003847.

40. Sarid R, Flore O, Bohenzky RA, Chang Y, Moore PS. 1998. Transcription Mapping of the Kaposi’s Sarcoma-Associated Herpesvirus (Human Herpesvirus 8) Genome in a Body Cavity-Based Lymphoma Cell Line (BC-1). J Virol https://doi.org/10.1128/jvi.72.2.1005-1012.1998.

41. Dittmer D, Lagunoff M, Renne R, Staskus K, Haase A, Ganem D. 1998. A Cluster of Latently Expressed Genes in Kaposi’s Sarcoma-Associated Herpesvirus. J Virol https://doi.org/10.1128/jvi.72.10.8309-8315.1998.

42. Schulz TF, Cesarman E. 2015. Kaposi Sarcoma-associated Herpesvirus: Mechanisms of oncogenesis. Curr Opin Virol.

43. Dittmer DP, Damania B. 2016. Kaposi sarcoma-associated herpesvirus: Immunobiology, oncogenesis, and therapy. J Clin Invest.

44. Rose TM, Bruce AG, Barcy S, Fitzgibbon M, Matsumoto LR, Ikoma M, Casper C, Orem J, Phipps W. 2018. Quantitative RNAseq analysis of Ugandan KS tumors reveals KSHV gene expression dominated by transcription from the LTd downstream latency promoter. PLoS Pathog https://doi.org/10.1371/journal.ppat.1007441.

45. Sadler R, Wu L, Forghani B, Renne R, Zhong W, Herndier B, Ganem D. 1999. A complex translational program generates multiple novel proteins from the latently expressed kaposin (K12) locus of Kaposi’s sarcoma-associated herpesvirus. J Virol.

46. McCormick C, Ganem D. 2006. Phosphorylation and Function of the Kaposin B Direct Repeats of Kaposi’s Sarcoma-Associated Herpesvirus. J Virol https://doi.org/10.1128/jvi.02331-05.

47. Kliche S, Nagel W, Kremmer E, Atzler C, Ege A, Knorr T, Koszinowski U, Kolanus W, Haas J. 2001. Signaling by human herpesvirus 8 kaposin a through direct membrane recruitment of cytohesin-1. Mol Cell https://doi.org/10.1016/S1097-2765(01)00227-1.

48. Muralidhar S, Pumfery AM, Hassani M, Sadaie MR, Kishishita M, Brady JN, Doniger J, Medveczky P, Rosenthal LJ. 1998. Identification of kaposin (open reading frame K12) as a human herpesvirus 8 (Kaposi’s sarcoma-associated herpesvirus) transforming gene. J Virol.

49. Tomkowicz B, Singh SP, Cartas M, Srinivasan A. 2002. Human herpesvirus-8 encoded kaposin: Subcellular localization using immunofluorescence and biochemical approaches. DNA Cell Biol https://doi.org/10.1089/10445490252925413.

50. Forte E, Raja AN, Shamulailatpam P, Manzano M, Schipma MJ, Casey JL, Gottwein E. 2015. MicroRNA-Mediated Transformation by the Kaposi’s Sarcoma-Associated Herpesvirus Kaposin Locus. J Virol https://doi.org/10.1128/jvi.03317-14.

51. Corcoran JA, Johnston BP, McCormick C. 2015. Viral Activation of MK2-hsp27-p115RhoGEF-RhoA Signaling Axis Causes Cytoskeletal Rearrangements, P-body Disruption and ARE-mRNA Stabilization. PLoS Pathog https://doi.org/10.1371/journal.ppat.1004597.

52. McCormick C, Ganem D. 2005. The kaposin B protein of KSHV activates the p38/MK2 pathway and stabilizes cytokine mRNAs. Science (80-) https://doi.org/10.1126/science.1105779.

53. Corcoran JA, Khaperskyy DA, Johnston BP, King CA, Cyr DP, Olsthoorn A V., McCormick C. 2012. Kaposi’s Sarcoma-Associated Herpesvirus G-Protein-Coupled Receptor Prevents AU-Rich-Element-Mediated mRNA Decay. J Virol https://doi.org/10.1128/jvi.00597-12.

54. Blanco FF, Sanduja S, Deane NG, Blackshear PJ, Dixon DA. 2014. Transforming Growth Factor Regulates P-Body Formation through Induction of the mRNA Decay Factor Tristetraprolin. Mol Cell Biol https://doi.org/10.1128/mcb.01020-13.

55. Franks TM, Lykke-Andersen J. 2007. TTP and BRF proteins nucleate processing body formation to silence mRNAs with AU-rich elements. Genes Dev https://doi.org/10.1101/gad.1494707.

56. Vindry C, Marnef A, Broomhead H, Twyffels L, Ozgur S, Stoecklin G, Llorian M, Smith CW, Mata J, Weil D, Standart N. 2017. Dual RNA Processing Roles of Pat1b via Cytoplasmic Lsm1-7 and Nuclear Lsm2-8 Complexes. Cell Rep https://doi.org/10.1016/j.celrep.2017.06.091.

57. Hubstenberger A, Courel M, Bénard M, Souquere S, Ernoult-Lange M, Chouaib R, Yi Z, Morlot JB, Munier A, Fradet M, Daunesse M, Bertrand E, Pierron G, Mozziconacci J, Kress M, Weil D. 2017. P-Body Purification Reveals the Condensation of Repressed mRNA Regulons. Mol Cell https://doi.org/10.1016/j.molcel.2017.09.003.

58. Brulois KF, Chang H, Lee AS-Y, Ensser A, Wong L-Y, Toth Z, Lee SH, Lee H-R, Myoung J, Ganem D, Oh T-K, Kim JF, Gao S-J, Jung JU. 2012. Construction and Manipulation of a New Kaposi’s Sarcoma-Associated Herpesvirus Bacterial Artificial Chromosome Clone. J Virol https://doi.org/10.1128/jvi.01019-12.

59. Myoung J, Ganem D. 2011. Generation of a doxycycline-inducible KSHV producer cell line of endothelial origin: Maintenance of tight latency with efficient reactivation upon induction. J Virol Methods https://doi.org/10.1016/j.jviromet.2011.03.012.

60. Kearse M, Moir R, Wilson A, Stones-Havas S, Cheung M, Sturrock S, Buxton S, Cooper A, Markowitz S, Duran C, Thierer T, Ashton B, Meintjes P, Drummond A. 2012. Geneious Basic: An integrated and extendable desktop software platform for the organization and analysis of sequence data. Bioinformatics https://doi.org/10.1093/bioinformatics/bts199.

61. Yoo SM, Ahn AK, Seo T, Hong HB, Chung MA, Jung SD, Cho H, Lee MS. 2008. Centrifugal enhancement of Kaposi’s sarcoma-associated virus infection of human endothelial cells in vitro. J Virol Methods https://doi.org/10.1016/j.jviromet.2008.07.026.

62. Carpenter AE, Jones TR, Lamprecht MR, Clarke C, Kang IH, Friman O, Guertin DA, Chang JH, Lindquist RA, Moffat J, Golland P, Sabatini DM. 2006. CellProfiler: Image analysis software for identifying and quantifying cell phenotypes. Genome Biol https://doi.org/10.1186/gb-2006-7-10-r100.

63. Cannon JS, Ciufo D, Hawkins AL, Griffin CA, Borowitz MJ, Hayward GS, Ambinder RF. 2000. A New Primary Effusion Lymphoma-Derived Cell Line Yields a Highly Infectious Kaposi’s Sarcoma Herpesvirus-Containing Supernatant. J Virol https://doi.org/10.1128/jvi.74.21.10187-10193.2000.

64. Tischer BK, Von Einem J, Kaufer B, Osterrieder N. 2006. Two-step Red-mediated recombination for versatile high-efficiency markerless DNA manipulation in Escherichia coli. Biotechniques https://doi.org/10.2144/000112096.

65. Santiago JC, Goldman JD, Zhao H, Pankow AP, Okuku F, Schmitt MW, Chen LH, Hill CA, Casper C, Phipps WT, Mullins JI. 2021. Intra-host changes in Kaposi sarcoma-associated herpesvirus genomes in Ugandan adults with Kaposi sarcoma. PLoS Pathog https://doi.org/10.1371/journal.ppat.1008594.

66. BeltCappellino A, Majerciak V, Lobanov A, Lack J, Cam M, Zheng Z-M. 2019. CRISPR/Cas9-Mediated Knockout and In Situ Inversion of the ORF57 Gene from All Copies of the Kaposi’s Sarcoma-Associated Herpesvirus Genome in BCBL-1 Cells. J Virol https://doi.org/10.1128/jvi.00628-19.

67. Sallah N, Palser AL, Watson SJ, Labo N, Asiki G, Marshall V, Newton R, Whitby D, Kellam P, Barroso I. 2018. Genome-wide sequence analysis of Kaposi sarcoma-associated herpesvirus shows diversification driven by recombination. J Infect Dis https://doi.org/10.1093/infdis/jiy427.

68. Tørresen OK, Star B, Mier P, Andrade-Navarro MA, Bateman A, Jarnot P, Gruca A, Grynberg M, Kajava A V., Promponas VJ, Anisimova M, Jakobsen KS, Linke D. 2019. Tandem repeats lead to sequence assembly errors and impose multi-level challenges for genome and protein databases. Nucleic Acids Res.

69. Jain V, Plaisance-Bonstaff K, Sangani R, Lanier C, Dolce A, Hu J, Brulois K, Haecker I, Turner P, Renne R, Krueger B. 2016. A toolbox for Herpesvirus miRNA research: Construction of a complete set of KSHV miRNA Deletion Mutants. Viruses https://doi.org/10.3390/V8020054.

70. Russo JJ, Bohenzky RA, Chien MC, Chen J, Yan M, Maddalena D, Parry JP, Peruzzi D, Edelman IS, Chang Y, Moore PS. 1996. Nucleotide sequence of the Kaposi sarcoma-associated herpesvirus (HHV8) Proceedings of the National Academy of Sciences of the United States of America.

71. Dheekollu J, Chen H-S, Kaye KM, Lieberman PM. 2013. Timeless-Dependent DNA Replication-Coupled Recombination Promotes Kaposi’s Sarcoma-Associated Herpesvirus Episome Maintenance and Terminal Repeat Stability. J Virol https://doi.org/10.1128/jvi.02211-12.

72. Deutsch E, Cohen A, Kazimirsky G, Dovrat S, Rubinfeld H, Brodie C, Sarid R. 2004. Role of Protein Kinase C δ in Reactivation of Kaposi’s Sarcoma-Associated Herpesvirus. J Virol https://doi.org/10.1128/jvi.78.18.10187-10192.2004.

73. Miller G, Rigsby MO, Heston L, Grogan E, Sun R, Metroka C, Levy JA, Gao S-J, Chang Y, Moore P. 1996. Antibodies to Butyrate-Inducible Antigens of Kaposi’s Sarcoma– Associated Herpesvirus in Patients with HIV-1 Infection. N Engl J Med https://doi.org/10.1056/nejm199605163342003.

74. Chen J, Ueda K, Sakakibara S, Okuno T, Parravicini C, Corbellino M, Yamanishi K. 2001. Activation of latent Kaposi’s sarcoma-associated herpesvirus by demethylation of the promoter of the lytic transactivator. Proc Natl Acad Sci U S A https://doi.org/10.1073/pnas.051004198.

75. Renne R, Zhong W, Herndier B, McGrath M, Abbey N, Kedes D, Ganem D. 1996. Lytic growth of Kaposi’s sarcoma-associated herpesvirus (human herpesvirus 8) in culture. Nat Med https://doi.org/10.1038/nm0396-342.

76. Ellison TJ, Kedes DH. 2014. Variable episomal silencing of a recombinant herpesvirus renders its encoded GFP an unreliable marker of infection in primary cells. PLoS One https://doi.org/10.1371/journal.pone.0111502.

77. Haddad CO, Kalt I, Shovman Y, Xia L, Schlesinger Y, Sarid R, Parnas O. 2021. Targeting the Kaposi’s sarcoma-associated herpesvirus genome with the CRISPR-Cas9 platform in latently infected cells. Virol J https://doi.org/10.1186/s12985-021-01527-x.

78. Wen J, Zhao S, He D, Yang Y, Li Y, Zhu S. 2012. Preparation and characterization of egg yolk immunoglobulin Y specific to influenza B virus. Antiviral Res https://doi.org/10.1016/j.antiviral.2011.11.005.

79. Brulois K, Toth Z, Wong L-Y, Feng P, Gao S-J, Ensser A, Jung JU. 2014. Kaposi’s Sarcoma-Associated Herpesvirus K3 and K5 Ubiquitin E3 Ligases Have Stage-Specific Immune Evasion Roles during Lytic Replication. J Virol https://doi.org/10.1128/jvi.00873-14.

80. Butnaru M, Gaglia MM. 2019. The kaposi’s sarcoma-associated herpesvirus protein orf42 is required for efficient virion production and expression of viral proteins. Viruses https://doi.org/10.3390/v11080711.

81. Jiang HY, Yang WH, Gulick T, Bloch KD, Bloch DB. 2005. Ge-1 is a central component of the mammalian cytoplasmic mRNA processing body. RNA https://doi.org/10.1261/rna.2142405.

82. Standart N, Weil D. 2018. P-Bodies: Cytosolic Droplets for Coordinated mRNA Storage. Trends Genet.

83. Tenekeci U, Poppe M, Beuerlein K, Buro C, Müller H, Weiser H, Kettner-Buhrow D, Porada K, Newel D, Xu M, Chen ZJ, Busch J, Schmitz ML, Kracht M. 2016. K63-Ubiquitylation and TRAF6 Pathways Regulate Mammalian P-Body Formation and mRNA Decapping. Mol Cell https://doi.org/10.1016/j.molcel.2016.05.017.

84. Seto E, Yoshida-Sugitani R, Kobayashi T, Toyama-Sorimachi N. 2015. The assembly of EDC4 and Dcp1a into processing bodies is critical for the translational regulation of IL-6. PLoS One https://doi.org/10.1371/journal.pone.0123223.

85. Robinson C-A, Singh GK, Castle EL, Boudreau BQ, Corcoran JA. 2021. The NDP52/CALCOCO2 selective autophagy receptor controls processing body disassembly. bioRxiv.

86. Ciufo DM, Cannon JS, Poole LJ, Wu FY, Murray P, Ambinder RF, Hayward GS. 2001. Spindle Cell Conversion by Kaposi’s Sarcoma-Associated Herpesvirus: Formation of Colonies and Plaques with Mixed Lytic and Latent Gene Expression in Infected Primary Dermal Microvascular Endothelial Cell Cultures. J Virol https://doi.org/10.1128/jvi.75.12.5614-5626.2001.

87. Grossmann C, Podgrabinska S, Skobe M, Ganem D. 2006. Activation of NF-κB by the Latent vFLIP Gene of Kaposi’s Sarcoma-Associated Herpesvirus Is Required for the Spindle Shape of Virus-Infected Endothelial Cells and Contributes to Their Proinflammatory Phenotype. J Virol https://doi.org/10.1128/jvi.01603-05.

88. Alkharsah KR, Singh V V., Bosco R, Santag S, Grundhoff A, Konrad A, Sturzl M, Wirth D, Dittrich-Breiholz O, Kracht M, Schulz TF. 2011. Deletion of Kaposi’s Sarcoma-Associated Herpesvirus FLICE Inhibitory Protein, vFLIP, from the Viral Genome Compromises the Activation of STAT1-Responsive Cellular Genes and Spindle Cell Formation in Endothelial Cells. J Virol https://doi.org/10.1128/jvi.00226-11.

89. Yoo J, Kang J, Lee HN, Aguilar B, Kafka D, Lee S, Choi I, Lee J, Ramu S, Haas J, Koh CJ, Hong YK. 2010. Kaposin-B enhances the PROX1 mRNA stability during lymphatic reprogramming of vascular endothelial cells by Kaposi’s sarcoma herpes virus. PLoS Pathog https://doi.org/10.1371/journal.ppat.1001046.

90. Chang HC, Hsieh TH, Lee YW, Tsai CF, Tsai YN, Cheng CC, Wang HW. 2016. c-Myc and viral cofactor Kaposin B co-operate to elicit angiogenesis through modulating miRNome traits of endothelial cells. BMC Syst Biol https://doi.org/10.1186/s12918-015-0242-3.

91. Aguilar B, Choi I, Choi D, Chung HK, Lee S, Yoo J, Lee YS, Maeng YS, Lee HN, Park E, Kim KE, Kim NY, Baik JM, Jung JU, Koh CJ, Hong YK. 2012. Lymphatic reprogramming by Kaposi sarcoma herpes virus promotes the oncogenic activity of the virus-encoded G-protein-coupled receptor. Cancer Res https://doi.org/10.1158/0008-5472.CAN-12-1229.

92. Toth Z, Papp B, Brulois K, Choi YJ, Gao SJ, Jung JU. 2016. LANA-Mediated Recruitment of Host Polycomb Repressive Complexes onto the KSHV Genome during De Novo Infection. PLoS Pathog https://doi.org/10.1371/journal.ppat.1005878.

93. Gallo A, Lampe M, Günther T, Brune W. 2017. The Viral Bcl-2 Homologs of Kaposi’s Sarcoma-Associated Herpesvirus and Rhesus Rhadinovirus Share an Essential Role for Viral Replication. J Virol https://doi.org/10.1128/jvi.01875-16.

94. Campbell M, Watanabe T, Nakano K, Davis RR, Lyu Y, Tepper CG, Durbin-Johnson B, Fujimuro M, Izumiya Y. 2018. KSHV episomes reveal dynamic chromatin loop formation with domain-specific gene regulation. Nat Commun https://doi.org/10.1038/s41467-017-02089-9.

95. West JA, Wicks M, Gregory SM, Chugh P, Jacobs SR, Zhang Z, Host KM, Dittmer DP, Damania B. 2014. An Important Role for Mitochondrial Antiviral Signaling Protein in the Kaposi’s Sarcoma-Associated Herpesvirus Life Cycle. J Virol https://doi.org/10.1128/jvi.03226-13.

96. Barbera AJ, Chodaparambil J V., Kelley-Clarke B, Joukov V, Walter JC, Luger K, Kaye KM. 2006. The nucleosomal surface as a docking station for Kaposi’s sarcoma herpesvirus LANA. Science (80-) https://doi.org/10.1126/science.1120541.

97. Hellert J, Weidner-Glunde M, Krausze J, Richter U, Adler H, Fedorov R, Pietrek M, Rückert J, Ritter C, Schulz TF, Lührs T. 2013. A Structural Basis for BRD2/4-Mediated Host Chromatin Interaction and Oligomer Assembly of Kaposi Sarcoma-Associated Herpesvirus and Murine Gammaherpesvirus LANA Proteins. PLoS Pathog https://doi.org/10.1371/journal.ppat.1003640.

98. Vladimirova O, de Leo A, Deng Z, Wiedmer A, Hayden J, Lieberman PM. 2021. Phase separation and DAXX redistribution contribute to LANA nuclear body and KSHV genome dynamics during latency and reactivation. PLoS Pathog https://doi.org/10.1371/journal.ppat.1009231.

99. Lin CL, Li H, Wang Y, Zhu FX, Kudchodkar S, Yuan Y. 2003. Kaposi’s Sarcoma-Associated Herpesvirus Lytic Origin (ori-Lyt)-Dependent DNA Replication: Identification of the ori-Lyt and Association of K8 bZip Protein with the Origin. J Virol https://doi.org/10.1128/jvi.77.10.5578-5588.2003.

100. Wang Y, Li H, Chan MY, Zhu FX, Lukac DM, Yuan Y. 2004. Kaposi’s Sarcoma-Associated Herpesvirus ori-Lyt-Dependent DNA Replication: cis-Acting Requirements for Replication and ori-Lyt-Associated RNA Transcription. J Virol https://doi.org/10.1128/jvi.78.16.8615-8629.2004.

101. Wang Y, Tang Q, Maul GG, Yuan Y. 2006. Kaposi’s Sarcoma-Associated Herpesvirus ori-Lyt-Dependent DNA Replication: DualRole of Replication and Transcription Activator. J Virol https://doi.org/10.1128/jvi.00990-06.

102. Liu D, Wang Y, Yuan Y. 2018. Kaposi’s Sarcoma-Associated Herpesvirus K8 Is an RNA Binding Protein That Regulates Viral DNA Replication in Coordination with a Noncoding RNA. J Virol https://doi.org/10.1128/jvi.02177-17.

103. Sharma NR, Majerciak V, Kruhlak MJ, Yu L, Kang JG, Yang A, Gu S, Fritzler MJ, Zheng ZM. 2019. KSHV RNA-binding protein ORF57 inhibits P-body formation to promote viral multiplication by interaction with Ago2 and GW182. Nucleic Acids Res https://doi.org/10.1093/nar/gkz683.

104. Wang X, Chang L, Wang H, Su A, Wu Z. 2017. Dcp1a and GW182 Induce Distinct Cellular Aggregates and Have Different Effects on microRNA Pathway. DNA Cell Biol https://doi.org/10.1089/dna.2017.3633.

105. Johnston BP, Pringle ES, McCormick C. 2019. KSHV activates unfolded protein response sensors but suppresses downstream transcriptional responses to support lytic replication. PLoS Pathog https://doi.org/10.1371/journal.ppat.1008185.

